# Mapping the Ligand-dependent Remodeling of the Conformational Entropy Landscape in Neurotensin Receptor 1 by NMR-guided Molecular Simulations

**DOI:** 10.64898/2026.01.17.700126

**Authors:** Scott A. Robson, Fabian Bumbak, Supriyo Bhattacharya, Wijnand J. C. van der Velden, Nagarajan Vaidehi, Joshua J. Ziarek

## Abstract

This study presents a comprehensive analysis of the dynamic properties and allosteric regulation mechanisms of Class A G protein-coupled receptors (GPCRs) by integrating molecular dynamics (MD) simulations with nuclear magnetic resonance (NMR) relaxation measurements. Utilizing generalized order parameters derived from NMR data and MD trajectories, we quantitatively assess conformational entropy changes that occur during receptor activation and ligand binding events. This approach enables a detailed characterization of protein flexibility at multiple timescales, revealing how dynamic fluctuations contribute to allosteric signal transmission within the receptor. Our results demonstrate that conformational entropy plays a pivotal role in modulating the functional states of Class A GPCRs, influencing the equilibrium between inactive and active conformations. By elucidating the interplay between structural dynamics and allostery, this work advances the molecular-level understanding of GPCR function and highlights the importance of entropy-driven effects in receptor signaling. The integrative methodology and findings provide a valuable framework for future investigations aimed at targeting receptor dynamics in drug discovery and rational design of allosteric modulators.

## INTRODUCTION

G protein-coupled receptors (GPCRs) constitute the largest superfamily of membrane proteins in the human genome and play a pivotal role in regulating diverse physiological processes by transducing extracellular signals into intracellular responses. Their extensive presence and accessibility have rendered them the most thoroughly investigated class of drug targets to date. As of 2017, they accounted for approximately 34% (475) of all therapeutics approved by the US Food and Drug Administration (FDA), targeting over 100 distinct GPCRs^1^.

Traditionally, drug discovery has relied predominantly on structure-based drug design, employing high-resolution structures derived from X-ray crystallography and cryo-electron microscopy^2^. While these static models have been instrumental in optimizing ligand affinity, they frequently fall short of accurately predicting efficacy and selectivity because of mechanisms such as partial and biased agonism^3,4^. This limitation arises because static structures typically capture low-energy activation endpoints, such as a fully activated state when bound to a transducer^5,6^ or an inactive state^7^ stabilized by an inverse agonist^8,9^. These endpoints obscure transitional or high-energy states, thereby skewing drug discovery towards states that static-structure methodologies can capture. GPCRs are increasingly recognized as dynamic conformational ensembles in which ligand binding regulates function by shifting the equilibrium populations of several possible substates{Kobilka, 2007 #393} {Motlagh, 2014 #286}, rather than as a binary switch. Consequently, the determinants of ligand efficacy are not solely encoded in an energy-minimized version of the receptor or its mean atomic coordinates. Instead, the thermodynamics of the ensemble play a significant role^3,11^.

This ensemble perspective requires a reevaluation of allosteric regulation, moving beyond traditional mechanical models that focus on specific structural changes. Cooper and Dryden proposed that allosteric communication can occur through changes in the frequency and amplitude of thermal fluctuations (entropy) in the absence of significant structural changes^12^. This dynamic allostery suggests that the entropic landscape is a critical determinant of ligand affinity and its ultimate function. Notably, the conformational entropy arising from fast picosecond-to-nanosecond (ps-ns) side-chain motions is a significant although not singular component of the free energy of molecular-binding events^13^. Nuclear magnetic resonance (NMR) relaxation studies have demonstrated that these fast motions, particularly those of methyl-bearing side chains, can serve as a robust dynamic proxy for conformational entropy, revealing that entropy can be a dominant driver of high-affinity molecular recognition^14,15^.

Neurotensin receptor 1 (NTS1), a prototypical peptide-binding class A GPCR, serves as a model system for dissecting the thermodynamic drivers of ligand binding. Neurotensin and its high-affinity receptor NTS1 are implicated in multiple physiological functions, including feeding, energy balance, pain modulation, thermoregulation, stress, and cardiovascular regulation^16^. NTS1 also modulates dopaminergic neurotransmission and is a validated therapeutic target for schizophrenia, drug addiction, and cancer^17^. Recent structural studies have revealed that agonists, such as the C-terminal hexapeptide of neurotensin (NT) NT8-13, select the contracted form of the orthosteric binding pocket of the receptor^8,17^. Conversely, inverse agonists (IvAs), such as SR142948A, shift the ensemble away from contraction and select an expanded pocket with a distinct helical arrangement^8^. Although these structures define the spatial boundaries of the orthosteric site, recent evidence using chemical shift-based NMR metrics, such as S_MCS,_ suggests that various ligands can selectively tune the global rigidity of NTS1^18^. Specifically, the inverse agonist SR142948A appears to act as an entropic clamp, rigidifying the receptor relative to the flexible apo and agonist-bound states. Furthermore, the discovery of biased allosteric modulators (BAMs), such as ML314^19^ and SBI-553^6^, which promote arrestin recruitment while dampening G protein activation, underscores the need to understand how ligands sculpt the dynamic landscape of receptors to achieve functional selectivity.

The rigorous quantification of methyl side-chain order parameters (*O*^*2*^_*axis*_) in GPCRs has been historically impeded by stringent isotope labeling requirements, specifically uniform deuteration. Notably, Clark et al. overcame this barrier using a *Pichia pastoris* expression system to generate highly deuterated Adenosine A_2A_ receptor (A_2A_R) samples, enabling the application of proton triple-quantum (3Q) relaxation experiments. This study provided the first direct evidence that the inverse agonist ZM241385 suppresses fast ps-ns motions at the G protein-binding interface relative to the agonist-bound state^20^. Similarly, Baumann et al. applied this 3Q methodology to a thermostabilized *α*_1B_-adrenergic receptor produced in *E. coli*, revealing that the peptide ligand ρ-TIA allosterically modulates the flexibility of specific isoleucine residues near the activation microswitches^21^. Complementing these studies are investigations utilizing alternative relaxation strategies in non-deuterated eukaryotic backgrounds, such as the ^13^C-relaxation analysis of the β_2_-adrenergic receptor by Kofuku et al.^22^ and the conformational dynamics studies of the muscarinic M2 receptor by Xu et al.^23^, which have collectively established that ligand efficacy is intimately coupled to the redistribution of the receptor’s dynamic landscape.

To address the role of fast dynamic changes (ps to ns) in NTS1 upon ligand binding, we targeted methionine residues distributed throughout the NTS1 transmembrane domain to serve as site-specific reporters of local dynamics in a minimal methionine variant *enNTS1ΔM4* (Figure 1A) as previously reported^24^. The amplitude of motion for the methyl symmetry axis, defined geometrically using the S_d_–C_e_ bond vector, was quantified by the squared generalized order parameter (*O*^*2*^_*axis*_)^25,26^. As illustrated in Figure 1B, this parameter is determined by the fluctuations of the bond vector polar angles (θ,ϕ) and is intimately linked to the population distribution of the side-chain dihedral angle (*χ*_3_) among canonical *trans* and *+/-gauche* rotamers^27^. NMR-derived methyl side-chain dynamics in proteins have been observed to cluster into distinct motional classes (Figure 1C)^14^. These range from the highly flexible J’-class and J-class (typically *O*^*2*^_*axis*_ ≈0.25 and ≈0.35, respectively), which arise from rapid averaging between multiple rotameric wells, to the *α*-class (*O*^*2*^_*axis*_ ≈0.6) representing restricted rotameric jumps, and finally the *ω*-class (*O* ^*2*^_*axis*_ >0.8) characteristic of restricted librational motion within a single potential energy well^13,27,28^.

**Figure 1:**
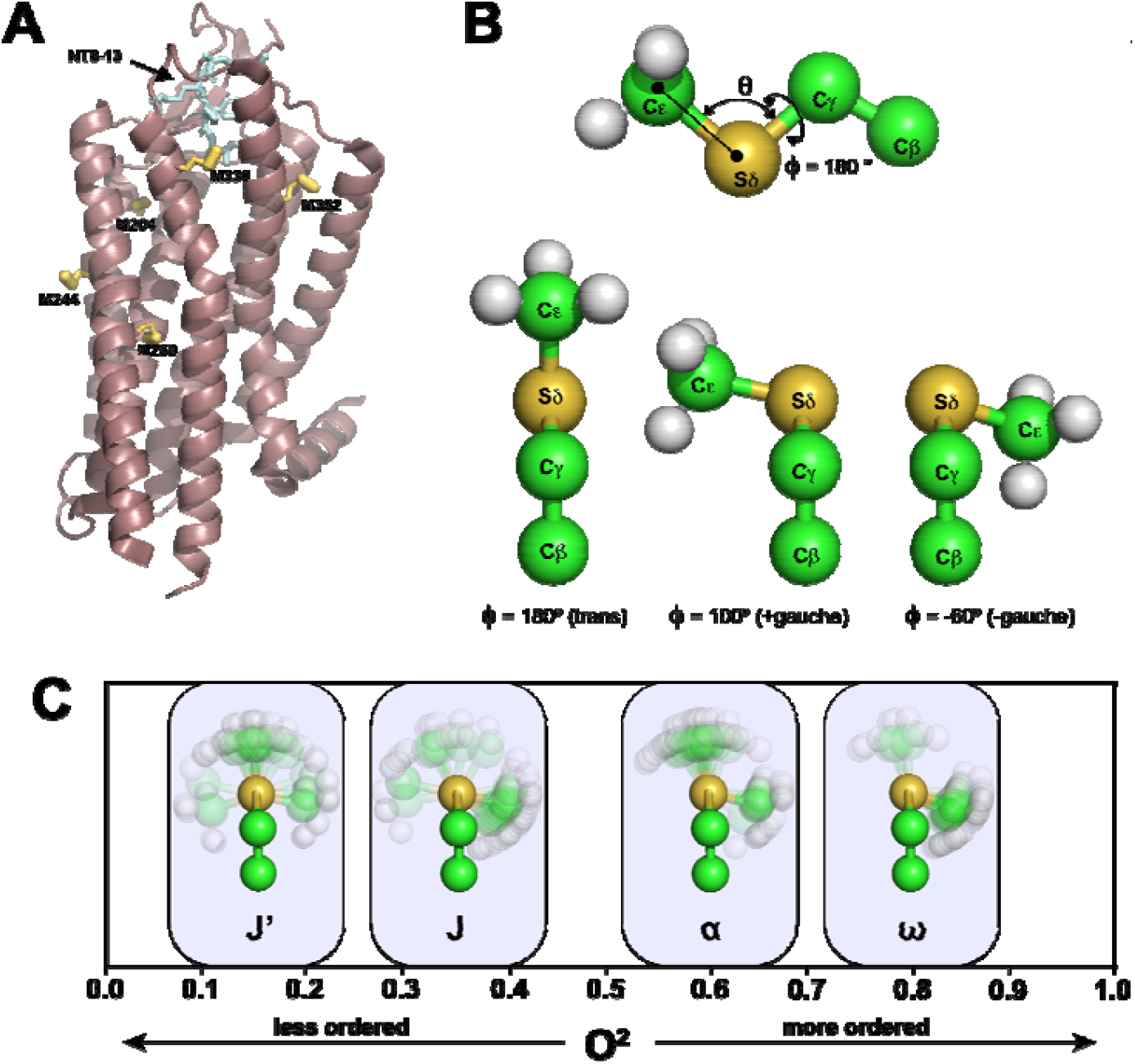
Positions and Behavior of Methionine Methyl Groups in Neurotensin Receptor 1 (NTS1). A) Cartoon representation of the thermostabilized variant of the rat neurotensin receptor 1 (rNTS1) (mauve) with NT8-13 bound (cyan), modelled from the PDB file 4BWB.pdb. The locations of the methionine residues present in our sample are indicated in yellow, with S*δ*-C_*ϵ*_ bonds indicated by thick lines. These methionines correspond to the minimal methionine construct (*enNTSM*Δ4) used in the present study. B) Model of methionine indicating the location of the θ and ϕ angles used to define the orientation of the O^2^_axis_ vector (top). The methyl orientations for the +/-*gauche* and *trans* positions are also shown. C) Distribution of Methionine O^2^_axis_, commonly found in proteins in solution, along with their conventional labelling according to the definitions in O’Brien *et. al*.^28^, namely J’, J, *α*, ω.

To rigorously quantify the thermodynamic contribution of these motions, entropic values from molecular ensembles in membrane-mimetic solutions are important. In this study, we aimed to bridge the gap between sparse experimental (NMR) observables and atomic-resolution thermodynamics to investigate the rapid changes in motion associated with the binding of the agonist (NT8-13) and inverse agonist (IvA) SR142948A to NTS1. To do so, we employed solution NMR triple quantum (3Q) spin relaxation of methionine methyl groups in a minimal methionine mutant of NTS1, *enNTS1ΔM4* to extract NMR based generalized order parameters (*O*^*2*^_*axis,NMR*_), which, as described above, directly report on the amplitude of ps-ns timescale fluctuations^29^. *The O*^*2*^_*axis,NMR*_ values were estimated from the experiment for the Apo (APO), Agonist (NT8-13), and Inverse Agonist (IvA) bound states of the NTS1 receptor in DDM micelles. Because of the limited *in vivo* access to atomic-level probes of dynamics in solution, especially at high molecular weight, we integrated our experimentally derived *O*^*2*^_*axis,NMR*_ values with *in silico* molecular dynamics (MD) simulations. By applying a modified maximum entropy reweighting algorithm^30^, we refined (reweighted) the MD frames to align the calculated *O*^*2*^_*axis,MD*_ values from the MD trajectories with the experimental NMR-based *O*^*2*^_*axis,NMR*_ values. This integrative methodology enables a detailed, quantitative characterization of side-chain conformational entropy changes upon ligand binding, providing new insights into the dynamic mechanisms underlying receptor activation and allosteric modulation.

## METHODS

### Design of *enNTS1ΔM4*

A “minimal-methionine” variant of enNTS1 (*enNTS1ΔM4*) was used, derived by site-directed mutagenesis of four solvent-exposed methionines (M181L, M267L, M293L, M408V) to reduce spectral overlap, retaining six endogenous methionines as NMR probes distributed across key receptor domains (orthosteric binding site, connector region/PIF motif, and extracellular helix termini). Functional validation assays confirmed ligand binding and coupling to G proteins and β-arrestins, supporting the construct’s physiological relevance. Resonance assignments were made via a “knock-in” mutagenesis strategy. Detailed construct design and validation are described in Bumbak et al.^31^.

### Expression of Isotopically Labeled *enNTS1ΔM4*

The thermostabilized neurotensin receptor 1 variant was expressed in *Escherichia coli* C43(DE3) as fusion proteins with N-terminal maltose-binding protein (MBP) and C-terminal monomeric ultra-stable green fluorescent protein (muGFP). For methyl-TROSY NMR studies, a methionine biosynthesis inhibition protocol was employed to incorporate [^13^CH_3_]-methionine selectively into a protonated background. Cultures were grown in defined medium with supplements and, at OD_600_ of 0.4, supplemented with isotopically labeled methionine and inhibitory amino acids to suppress endogenous methionine synthesis and isotopic scrambling. Detailed expression and labeling protocols are described in Bumbak et al.^31^.

### Preparation of NMR Samples

The NMR samples were prepared by exchanging the buffer solvent to 100% D_2_O to eliminate solvent signals and reduce proton-proton relaxation pathways. This exchange was performed through repeated concentration and dilution steps or size-exclusion chromatography (SEC) using D_2_O-based buffers. The final receptor concentration was adjusted to approximately 60–70 µM. Ligands were solubilized and added to the receptor samples as follows: the peptide agonist NT8-13 was prepared as 5–10 mM stock solutions in 100% D_2_O and added at a saturating concentration of 500 µM. The inverse agonist SR142948A was prepared as a 20 mM stock solution in 100% D_2_O; unlike many small molecules that require organic solvents, SR142948A was solubilized directly in deuterated water for these experiments.

### NMR Data Collection

NMR experiments were conducted on a Bruker Avance Neo spectrometer operating at 600 MHz. Methyl side-chain dynamics were probed using the triple-quantum (3Q) relaxation-based coherence transfer pulse sequence described by Sun et al.^29^. The experiment quantifies the buildup of “forbidden” 3Q coherences relative to the decay of “allowed” single-quantum (SQ) coherences to extract methyl axis order parameters.

The 3Q and SQ spectra were acquired in an interleaved manner to minimize artifacts arising from sample changes during data collection. To compensate for the inherently lower sensitivity of the forbidden coherence transfer pathway, the 3Q state was recorded with 936 scans, while the reference SQ state was recorded with 144 scans. The experiments employed a variable relaxation delay (T) sampled at seven time points: 0.4, 1.6, 3.0, 4.0, 6.0, 8.0, and 10.0 ms. Thus, a total of 14 spectra were acquired.

Spectral windows were optimized for the methyl region, with the ^1^H dimension set to a spectral width of 3571.4 Hz centered at 1.8 ppm, and the ^13^C dimension set to a spectral width of 678.7 Hz centered at 18.75 ppm. Data were acquired using non-uniform sampling (NUS) with a sampling density of approximately 28% (sampling 9 out of 32 complex points in the indirect dimension). Schedules were randomly and individually selected for each of the 14 spectra using the Poisson Gap method^32^. The NUS data were reconstructed using the hmsIST^32^. Spectral processing, including Fourier transformation, phasing, and apodization, was performed using NMRPipe^33^. The first delay time spectra of the SQ data are shown in Figure S1.

### Extraction of *η* and *δ* values from 3Q Relaxation Data

In traditional analysis of 3Q relaxation data, *η* (and *δ*) values are determined by calculating the ratio of 3Q to SQ signal intensities (I_3Q_ and I_SQ_) and fitting these values to Eq. 1 using non-linear least squares (NLS). Here, *η* represents the cross-correlated relaxation rate between the ^1^H nuclei and the ^13^C nucleus in methyl groups and is related to O^2^_axis_, while *δ* accounts for the relaxation rate due to ^1^H-^1^H dipolar interactions between the methyl protons and other nearby protons. Although sample deuteration can minimize these interactions, they cannot be eliminated.

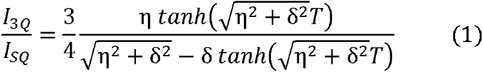

However, we encountered a significant problem using Eq. 1 with our data (See Figure S2): some fittings performed very poorly, yielding large errors or instability/convergence issues. Principally, the challenges were low signal-to-noise due to low sample concentration and high molecular weight, compounded by high levels of artifact noise from the high lipid signal concentration. The low signal-to-noise also meant that only a limited number of data points could be acquired before sample degradation. Thus, it was a challenge to use Eq. 1 with high noise and limited data. We reposed Eq. 1 as two simultaneous equations to fit the 3Q (numerator of Eq. 1) and SQ (denominator of Eq. 1) data independently, while sharing parameters. This approach increases the number of data points 2-fold since we no longer construct a ratio of data points. The results of this method are discussed in the results.

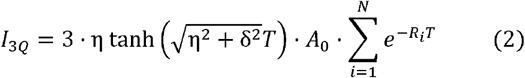

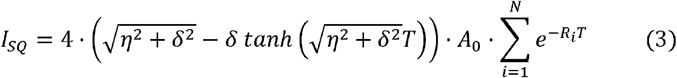

The extracted *η* values were converted to O^2^_axis_ estimates using the theory in Sun et. al.^29^ Eq. 4a shows how *η* is related to O^2^_axis_, while 4b is an algebraic rearrangement to express O^2^_axis_ in terms of *η*.

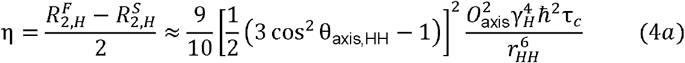

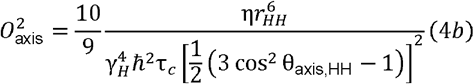

From Eq. 4b, O^2^_axis_ is a function of not only *η* but also physical and geometric constants (known) and the molecular correlation time, *τ*_c_. To estimate *τ*_c_, we used Small Angle Xray Scattering (SAXs) as detailed previously^24^. SEC-SAXS data for both Apo and NT8-13 bound forms showed similar scattering profiles (Figure S8). Guinier plots where used to estimate R_g_. Using the Stoke-Einstein relationship (Eq. 5) we estimated *τ*_c_, which permitted the estimation of O^2^axis.

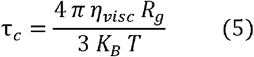

where *η*_*visc*_ is the viscosity of the solution. We estimated the viscosity of our solution from data obtained in Hardy et al^34^. We produced a third-order polynomial calibration curve (Figure S9) and estimated *η*_*visc*_ for D_2_O at 25 °C (1.099 cP).

Errors from SAXS estimation of *τ*_c_ and NMR estimation of *η* were propagated into O^2^_axis_ by modeling *τ*_c_ as a probabilistic variable in PyMC, calculated from a Gaussian distribution with a mean and standard deviation based on the value and error of R_g_ from SAXS.

### Molecular Dynamics Simulations

Molecular dynamics (MD) simulations were initiated from high-resolution crystal structures of the rat neurotensin receptor 1 (NTS1) in three distinct conformational states: the apo state (PDB: 6Z66)^8^, the agonist-bound state complexed with NT8-13 (PDB: 6YVR)^8^, and the inverse agonist-bound state complexed with SR142948A (PDB: 6Z4Q)^8^. Missing residues and side-chain atoms, as well as a micelle of 192 DDM molecules, were modeled, and the receptor systems were assembled using the CHARMM-GUI interface^35,36^. The receptor complexes were solvated in a cubic box using the TIP3P water model and neutralized with 0.15 M NaCl to mimic physiological ionic strength.

Simulations were performed using the CHARMM36 all-atom additive force field for proteins, lipids, and detergents^37^. Crucially, to ensure accurate reproduction of methyl side-chain dynamics required for NMR relaxation analysis, we utilized a modified version of the CHARMM36 force field as described by Hoffmann et al. ^38^. In this parameter set, the dihedral energy barriers (*V*_*dih*_) governing the rotation of methyl groups in alanine, methionine, threonine, valine, leucine, and isoleucine side chains were reparameterized against high-level coupled cluster [CCSD(T)] quantum chemical calculations to correct the systematic overestimation of rotational barriers present in the standard force field^38^. Ligand parameters for SR142948A and NT8-13 were generated using the CHARMM General Force Field (CGenFF).

Molecular dynamics simulations were performed using GROMACS (version 2021/2022) following the protocol described in Bower et al.^39^. Systems were energy-minimized, equilibrated with positional restraints gradually released, and production runs conducted in the NPT ensemble at 310 K and 1 bar. Temperature and pressure were controlled by the Nose-Hoover thermostat and Parrinello-Rahman barostat, respectively. Long-range electrostatics employed the Particle Mesh Ewald method with a 12 Å cutoff. Five independent 1-µs replicas were simulated for each conformational state (Apo, NT8-13, and IvA).

### O^2^_axis_ Calculations from MD Simulations

Prior to analysis, rotational and translational diffusion of the protein were removed from the production trajectories by least-squares fitting of the backbone C_*α*_ atoms to the initial crystal structure reference frame. The squared generalized order parameter was calculated via custom Python scripts implementing the analytical formalism described by Krishnan and Smith^27^. This approach quantifies the angular spatial restriction of the methyl symmetry axis (S_*δ*_-C_*ε*_ bond vector) within the molecular frame.

For every frame of the trajectory, the instantaneous bond vector *v(t)* was defined using the Cartesian coordinates of the sulfur (S_*δ*_) and methyl carbon (C_*ε*_) atoms and then normalized to 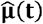:

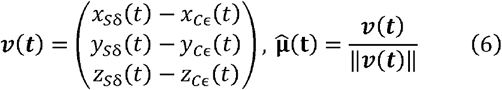

Alternatively, the orientation was parameterized in a local spherical coordinate system defined by the polar angle *θ* (the angle between the C_*γ*_ - S_*δ*_ and S_*δ*_ - C_*ε*_ bonds, represented as π - *θ* in the vector projection) and the azimuthal angle *ϕ* (the C_β_ - C_*γ*_ - S_*δ*_ - C_*ε*_ dihedral angle):

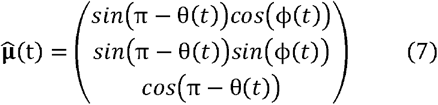

The squared order parameter O^2^_axis_, which ranges from 0 (isotropic disorder) to 1 (complete rigidity), was calculated from the ensemble averages of the Cartesian components 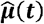 of using the expansion of the traceless second-rank tensor:

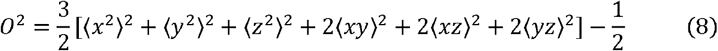

where angular brackets <…> denote the average over all snapshots in the trajectory.

To validate the ensemble-based calculations and inspect the decay of orientational memory, we computed the internal time correlation functions (TCFs) for the methionine S*δ*-C*ε* bond vectors as described by Hoffmann et al.^38^. TCFs were calculated using the MethylRelax code provided by Hoffmann and colleagues. The internal TCF, *C*_*int*_*(t)* describes the reorientation of the bond vector 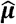 after removing overall tumbling:

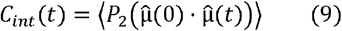

where *P*_2_ (*x*) = (3*x*^2^ − 1) is the second-order Legendre polynomial. The long-time limit of the internal TCF corresponds to the squared generalized order parameter, such that *O*^2^ = lim_*t*→ ∞_ *C*_*int* (*t*)_. This method assumes that the internal motions are statistically independent of the global molecular tumbling. The order parameters derived from the TCF plateau were compared against those calculated via the static tensor expansion to ensure consistency.

### Reweighting of MD Trajectories

To reconcile the molecular dynamics simulations with the experimental NMR data, we employed a Bayesian Maximum Entropy (BME) reweighting approach. Standard approaches assume observables are linear averages over the ensemble of the simulation (Bottaro et al., 2020). However, calculations of O^2^_axis_ (Eq. 9) depend non-linearly on ensemble weights due to the quadratic expansion of the averaged tensor components (Eq. 10).

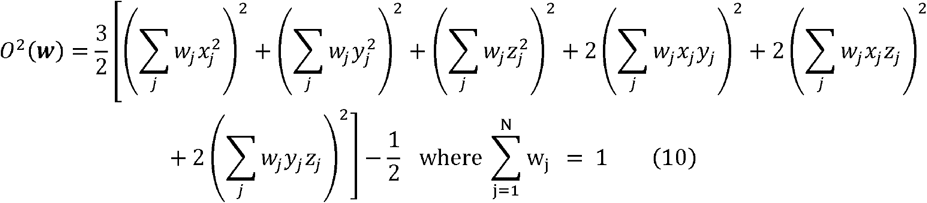

We wish to find weights (*w* = {*w*_1_, …*w*_*n*_} where 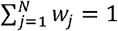) that minimize the objective function Eq. 11.

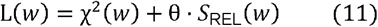

The discrepancy between the calculated and experimental order parameters was quantified by Eq. 12:

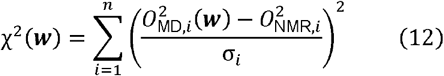

Where n is the number of O^2^_axis_ values from the NMR experiments. To prevent overfitting weights to the data, the deviation from initial simulation ensemble (the prior, denoted by 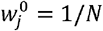) was penalized using the relative entropy of the Kullback-Leibler (KL) divergence in Eq. 13.

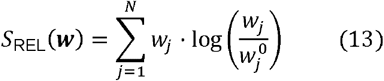

Due to the non-linearity of the function, the analytical Lagrange multiplier solution typically employed in Bayesian/MaxEnt reweighting^30,40,41^ was not applicable. Instead, the weights were determined through direct numerical minimization of the cost function *L*(*w*). The optimization was performed using the Limited-memory Broyden-Fletcher-Goldfarb-Shanno (L-BFGS-B) algorithm, utilizing analytically derived gradients of Eqs. 10, 12, and 13 with respect to weights. For algorithmic optimization, the weights were treated as unnormalized variables with box constraint (0 ≤ *w*_*u,j*_) to strictly enforce non-negativity. After convergence, the weights are normalized by the sum of unnormalized weights over the N frames. Optimizing the unnormalized constraints is considerably faster than optimizing the normalized weights with the additional constraint of 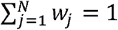. To speed up convergence with L-BFGS-B algorithm used in SciPy^42^, we used analytical gradients of Eq. 10, 12 and 13 which were derived with respect to the unnormalized weights *w*_*u,j*_.

The hyperparameter *θ* controls the trade-off between fitting the experimental data (low *θ*) and retaining the diversity of the original MD ensemble (high *θ*). An optimal value for *θ* was selected by performing the minimization over a logarithmic range of *θ* values and analyzing the resulting “L-curve” of *χ*^2^ versus *θ* and the effective fraction of frames (N_eff_), choosing a value at the elbow of the curve to avoid overfitting^30,43^.

### Calculation of Reweighted Conformational Entropy

To quantify the thermodynamic consequences of the reweighted ensembles, side-chain conformational entropies (*S*_*SC*_) were calculated using an information-theoretic histogram approach applied to the side-chain dihedral angles (*χ*). Unlike standard analyses that derive probabilities from raw population counts, we estimated the probability density functions of the dihedral angles using the optimized weights ( *w*_*j*_) obtained from the Maximum Entropy refinement. This ensures that the calculated entropies reflect the experimentally consistent ensemble rather than the prior force field’s raw sampling.

For a given dihedral angle, the continuous conformational space was discretized into 10° bins. This bin width was selected to capture both the rotameric jump contributions and the intra-well vibrational entropy, analogous to the fine-binning strategies described previously^44^. The reweighted probability of the k-th bin was calculated as the sum of the weights of all frames falling within that bin:

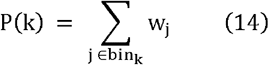

where ∑*P*(*k*) = 1. The conformational entropy for each residue was then computed using the Gibbs-Shannon formula:

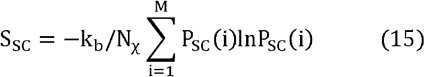

where *k*_*b*_ is the Boltzmann constant and N_χ_ is the number of *χ* angles in the residue, and M is the number of bins. Note, in this form, S_SC_ is dimensionless.

Accurate entropy estimation for long, flexible side-chains, particularly methionine residues containing the torsion, requires extensive sampling to ensure convergence of the dihedral probability distributions^44^. To assess convergence and estimate statistical uncertainty, the full trajectory (comprising 25,000 frames) was treated as one continuous segment of all frames. This was necessary due to the slow convergence of methionine residues in particular (Figure S15).

## RESULTS

### Measurement of O^2^_axis_ by NMR

^1^H-^13^C sofast HMQC spectra were collected on Apo, NT8-13, and IvA bound samples to confirm the resonance assignments previously reported by Bumbak et al.^18^. The HMQC spectrum for NT8-13 bound to *enNTS*1Δ*M*4 is shown in Figure 2A. Assignments of all systems analyzed, in all bound states, are shown in Figure S1. Subsequently, the 3Q relaxation experiment described by Sun et. al.^29^ was performed to estimate the *η* values. Figure 2B illustrates the time series data for both 3Q and single quantum (SQ) relaxation measurements.

**Figure 2:**
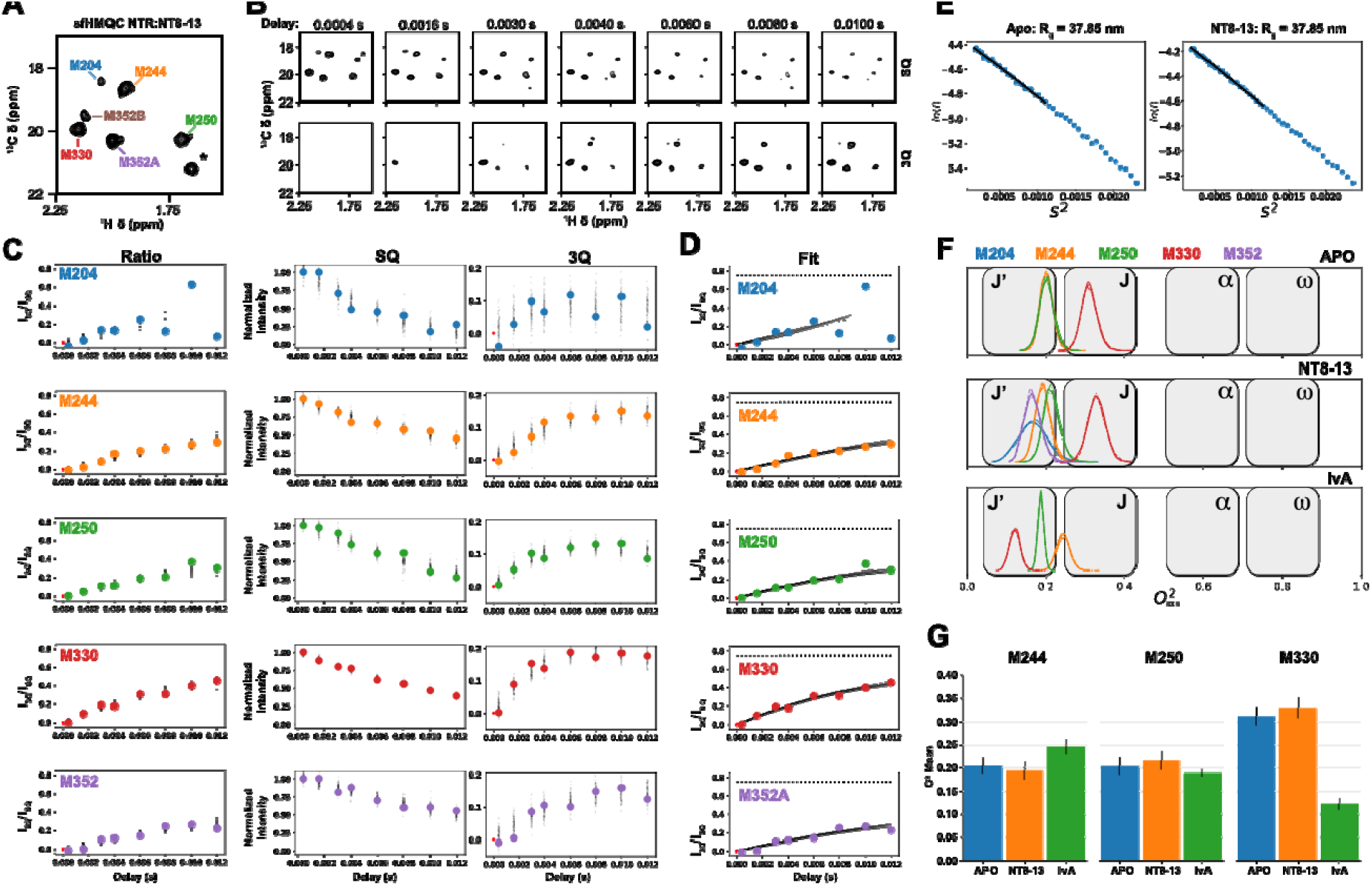
Estimation of O^2^_axis_ from peak identification, triple quantum relaxation experiments, model fitting, η estimation, R_g_ estimation and O^2^_**axis**_ calculation. A) sofast ^13^C HMQC of NT8-13 loaded *enTS*1Δ*M*4 with methionine assignments. (*) indicates an artifact peak resulting from a high detergent concentration. B) Delay time series spectra of the single-quantum (SQ) and triple-quantum (3Q) relaxation spectra. Again, (*) indicates a detergent peak that has shifted position due to a different ^13^C sweep width in the relaxation spectra compared to that in the HMQC. C) Simultaneous fitting of the data to the models described by Eq. 1 (ratio model) and Eq. 2 and 3 (SQ and 3Q models) with shared η and d parameters. Solid circles indicate the measured 3Q/SQ peak ratios (left column) and peak intensities normalized to 1.0 for the highest value in the data for each peak (center (SQ) and right columns (3Q)). Black dots indicate posterior predictive estimates of the data from the fitted models, including the fitted data noise terms for SQ and 3Q (see Methods). D) Distribution of 100 models (black lines) fit from posterior parameter values. Again, the filled circles indicate the ratio of the 3Q to the SQ peak intensities. E) Determination of the radius of gyration (R_g_) by SAXS. SEC-SAXS data for both the Apo and NT8-13-bound forms of NTR showed similar scattering profiles. The Guinier plots estimated R_g_ at 37.85 nm for both cases. F) Calculation of O^2^ from the distribution of ηin (C) and (D) and *τ*_*c*_ from Rg (E) using Eq. 4. The distribution of O^2^_axis_ for the identified peaks in all bound states examined is color-coded according to the legend across the top, with the bound state indicated in the top right of each plot. G) Bar plot of O^2^_axis_ versus bound state, separated by amino acid (M244, M250, M330).

However, we encountered a significant problem using Eq. 1 with our data (See Figure S1), with some fittings behaving very poorly, and fittings gave large errors or were unstable/non-converging. Principally, the challenges were low signal-to-noise due to low sample concentration and high molecular weight, compounded by high levels of artifact noise from the high lipid signal concentration. The low signal-to-noise also meant that only a limited number of data points could be acquired before sample degradation. Thus, it was a challenge to use Eq. 1 with high noise and limited data. Instead, we used Eq. 2 and Eq. 3, to define a model with *η* and *δ* values along with an estimation of noise, which accounts for the error of the fit. This requires the inclusion of relaxation terms to accurately model signal relaxation during the time delays. Note that these terms have shared parameters between Eq. 2 and Eq. 3 as well, while the ratio method eliminates them^29,45^. These fits were performed using a Bayesian Parameter Estimation (BPE) protocol in the probabilistic programming language, PyMC^46^. Validation of this procedure was performed using example data provided by Vitaly Tugarinov for the small Ubiquitin and large *α*7*α*7 proteins^29^. In our experience, we found that two relaxation terms (N = 2 in Eq. 2 and 3) were required for many systems from the Tugarinov data, so we used two terms for all our fits with NTS1 (See Figure S3 - Ubiquitin data and Figure S4 - *α*7*α*7 data). The correlation between the traditional nonlinear least squares (NLS) approach and the monophasic (N=1) and biphasic (N=2) approaches shows that the biphasic approach better matches the NLS approach on low-noise data (See Figure S5). This enabled robust extraction of relaxation parameters even under high noise levels and sparse data (Figure S6, compared to Figure S1).

Posterior predictive points (black dots) along with the ratio, SQ, and 3Q data (solid, colored circles) are shown in Figure 2C for each methionine assignment in the NT8-13 bound state. Posterior predictive points indicate the level of uncertainty in the fitted model and are constant when fitting individual SQ (center column) and 3Q (right column) time series. However, the error distribution increases with time delay for the ratio model (left column). This is expected, since dividing by increasingly smaller values over time (I_SQ_) with the same error increases the uncertainty of the quotient. This protocol for handling errors more accurately models the noise in ratio data than a standard non-linear least-squares approach. The final model fits are shown in Figure 2D as 100 simulations of the curve generated from Eq. 1 using sampled values of *η* and *δ* from their fitted distributions. The success of this approach is particularly apparent for the high noise and relatively poor fit seen for M204 in Figure 2D. In this case, the estimates for *η* and *δ* do not ‘blow up’ and permit an estimate to be made, even if the error is quite large. The fitting results for APO and IvA bound states are shown in Figure S7.

Figure 2E shows Guinier plots for SEC-SAXS data collected on the Apo and NT8-13 samples. Linear extrapolation of this data gave R_g_ values of 37.85 nm for both Apo and NT8-13 samples. Raw scattering data for APO and NT8-13 bound NTS1 are shown in Figure S8. The consistency between these two results suggests that Apo and NT8-13 do not differ significantly in size or multimeric state. Using Eq. 5 and a calibrated estimate of the viscosity of our solution (Figure S9), we estimated *τ*_c_ to be 65.17 ns with a standard deviation of 0.48 ns.

Figure 2F show our O^2^_axis_ estimates from Eq. 2 and 3 fall into the J’ and J states, which is expected of methionine methyls^28^. The *η* and *δ* estimates and errors for every methionine assigned in Apo, NT8-13, and IvA states are shown in Figure 2G. This analysis reveals that the inverse agonist-bound state of M330^6.57^ is characterized by elevated side-chain dynamics, providing a kinetic resolution to prior spectral interpretations. Bumbak et al.^18^ reported a significant (45%) increase in peak intensity upon binding the inverse agonist SR142948A, which was originally attributed to a reduction in basal motions away from the millisecond time scale, which cause broadened (weak) peaks in NMR spectra. However, our result suggests this spectral sharpening arises from a shift to higher dynamics away from the millisecond time scale rather than rigidification. In the APO state, M330^6.57^ likely undergoes intermediate-to-slow exchange on the microsecond-millisecond timescale, resulting in exchange-induced line broadening. It’s important to note that the 3Q relaxation experiment is insensitive to the line broadening time scale and it is only able to capture changes in fast (ps-ns) motions. The binding of SR142948A, which utilizes a bulky adamantyl moiety to sterically wedge apart the extracellular tips of TM6 and TM7, may effectively unlock the local environment of M330^6.57^, while rigidifying F331^6.58^. This structural expansion likely shifts the side chain of M330^6.57^ into a lower O^2^_axis regime_, thereby eliminating the exchange broadening contribution and resulting in the strong signal intensity observed in the HMQC. Conversely, our measurement of an increase for O^2^_axis_ for M330^6.57^ in the NT8-13 complex relative to Apo may reflect a genuine restriction of conformational space, consistent with the agonist-induced contraction of the orthosteric pocket and the stabilization of the N-terminus/ECL2 “lid” over the binding site (compare Figure 2G APO with Figure 2F NT8-13 for M330) although the effect appears to be slight and within error.

M244^5.45^, positioned on helical turn extracellular to the conserved P^5.50^, functions as a sensor for the conformational status of the PIF motif (P^5.50^/I^3.40^/F^6.44^) and reports on activation level through the rearrangement of a core aromatic network involving Y154^3.37^ and F248^5.498^. In the presence of the inverse agonist SR142948A, it has been suggested that M244^5.45^ is stabilized in a deshielded environment characterized by its proximity to the edge of the Y154^3.37^ aromatic ring, while F248^5.49^ is displaced outside the immediate interaction sphere^31^. This hypothesized structural rigidification suppresses basal conformational fluctuations and results in a reported increase in ^1^H-^13^C HMQC NMR signal relative to the APO state^31^. Conversely, binding of NT8-13 was reported to shift M244^5.45^ into an intermediate to slow exchange regime with a loss of signal^31^. Figure 2G (M244) shows a slight increase in O^2^_axis_ compared to APO, which is in line with stabilization of this residue. On the other hand, there is little change in O^2^_axis_ for M244^5.45^ in the NT8-13 bound state compared to APO. Given that intermediate-time-scale changes were previously observed, it seems these changes are confined to the intermediate time scale.

M250^5.51^ shows little change from APO to NT8-13 to IvA on the O^2^_axis_ at the fast time scale. Agonist binding has previously been associated with an increase in intermediate exchange for M250^5.51^, whereas IvA binding did not change it. Our relatively flat O^2^_axis_ response to ligands suggests these changes are not present on the fast time scale.

### Molecular Dynamics Simulations

Figure 3 presents a comprehensive analysis of molecular dynamics (MD) simulations focusing on methionine dynamics in the NT8-13-bound state. Panel A displays the time correlation functions (TCFs) for all methionines, using the bond-vector definition of the x, y, and z coordinates as described by Eq. 6 and calculated according to Eq. 9. This panel shows five individual trajectories (black) and their average (green). In contrast, panel B shows the same set of TCFs calculated using the spherical-coordinate (angle) approach of Eq. 7. Panel C is a bar graph comparing results from Eq. 8 and Eq. 9, using either vectors (Eq. 6) or angles (Eq. 7).

**Figure 3:**
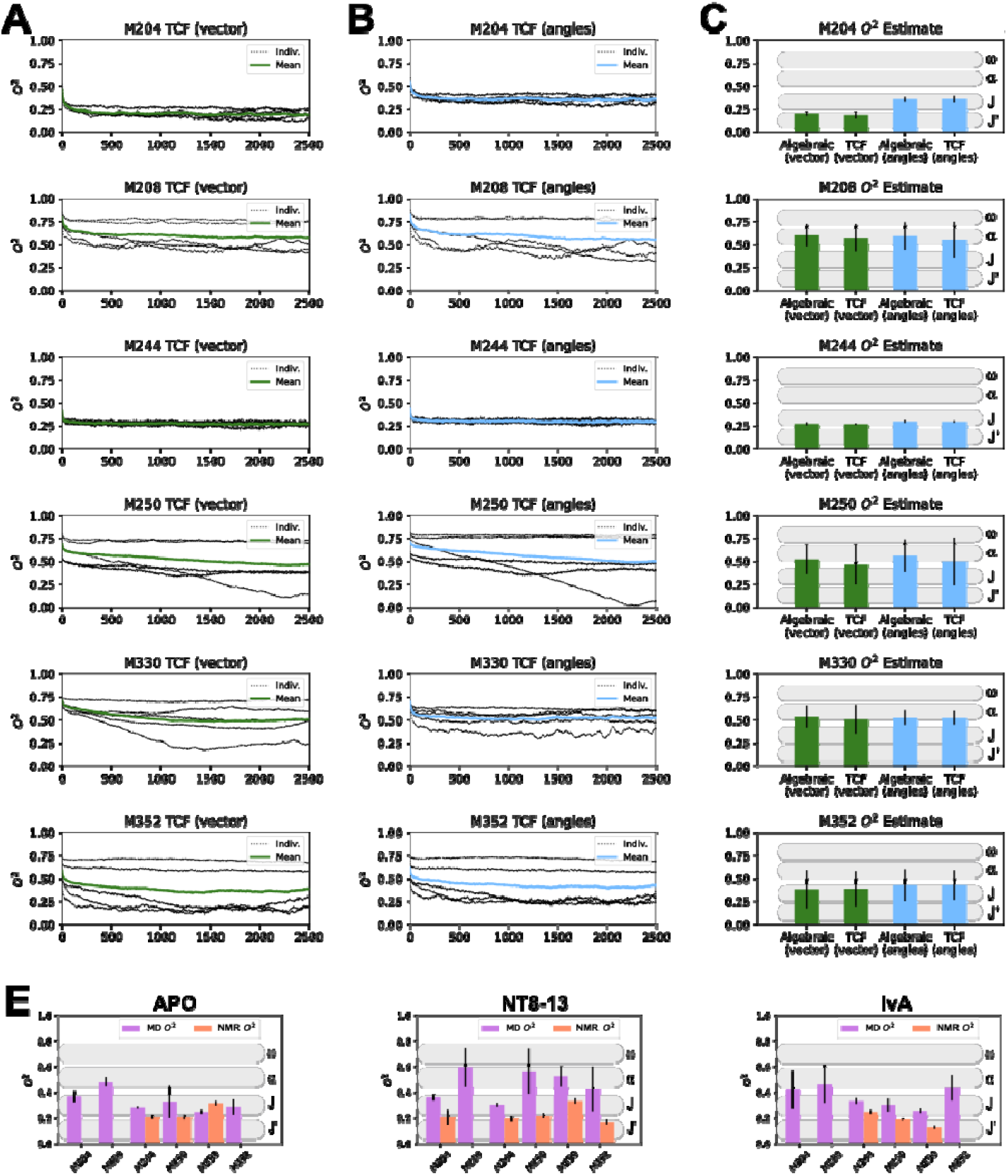
MD simulation analysis of *enNTS*1Δ*M*4 in the NT8-13 bound state. A) Time correlation function calculations for each methionine based on the normalized vector definition of the O^2^_axis_ bond (Eq. 5). The black dotted lines represent the individual MD simulation runs, with a solid green line indicating the average. B) Same as A) but using the spherical coordinate definition of the O^2^_axis_ bond (Eq. 6). and asolid cyan line for the average value. C) Summary bar graph of the average estimated O^2^_axis_values usingthe algebraic (Eq. 8) or last point values in the TCF (Eq. 9) for the vector definition (green) and angle definition (cyan) methods. Black vertical lines indicate the standard deviation of the average. D) Comparison of O^2^_axis,MD_ and O^2^_axis,NMR_. For Apo, NT8-13 and IvA bound states. Error bars indicate the standard deviation.

Generally, the O^2^_axis_ values are higher when using the angle approach of Eq. 7, most likely because this definition of the S_*δ*_-C_*ε*_ bond is more localized and is not subject to the minor molecular fluctuations that remain even after C_*α*_ alignment (see Methods). It is interesting to compare the heterogeneity of individual MD traces with the intermediate exchange observed in ^1^H-^13^C HMQC spectra^31^. For example, M250 in Figure 3 shows a rather inconsistent pattern in TCFs across 5 individual MD trajectories, suggesting that this residue can be sampled in either highly or lowly dynamic states. This could be consistent with this residue exhibiting intermediate exchange, as previously observed in ^1^H-^13^C HMQC spectra. The same can be said for M330 and M352 in Figure 3. In contrast, a shift into intermediate exchange has also been seen for M244 in ^1^H-^13^C HMQC spectra, however our MD results suggest this residue is consistently highly flexible.

Panel D integrates MD simulation results with experimental NMR data by comparing the order parameters derived from Eq. 8, where the x, y, and z coordinates are defined using Eq. 7. Generally, there is poor agreement between the O^2^_axis_ metrics, with MD O^2^_axis_ typically being higher than NMR O^2^_axis_. However, some consistencies are present. For example, in the IvA panel, O^2^_axis_ is consistently in the order M244 > M250 > M352 for both MD and NMR estimations.

### Bayesian/Maximum Entropy (BME) Reweighting Modification of MD Trajectories

While standard molecular dynamics simulations provided a high-resolution microscopic view of the NTS1 conformational landscape, quantitative analysis revealed discrepancies between the back-calculated O^2^_axis_ and the experimental NMR-derived values. Such deviations are frequently observed in atomistic simulations of proteins and are typically attributed to inherent imperfections in empirical force fields, particularly regarding methyl rotation barriers^30^ or the incomplete sampling of rare rotameric transitions that occur on timescales exceeding the trajectory length^38,44^.

To reconcile the microscopic details of the molecular dynamics simulations with the macroscopic experimental observables, we applied a Bayesian/Maximum Entropy (BME) reweighting algorithm to the initial MD ensembles^30^. As discussed in the Methods section, our approach directly determines the weights rather than using Lagrange multipliers, since the derived parameter O^2^_axis_ is a non-linear function of the weights. The regularization parameter *θ*, which balances the minimization of the error between calculated and experimental observables (*χ*^2^) against the maximization of the relative entropy (*S*_*REL*_), was determined using an L-curve analysis (Figure 4A). As *θ* decreases, the agreement with experimental data improves; however, this comes at the cost of reducing the effective ensemble size (*N*_*eff*_), potentially leading to overfitting where a small number of frames dominate the ensemble. We identified an optimal regime at *θ*=10(Figure 4A, dashed vertical line), located at the “elbow” of the *χ*^2^ versus *θ* curve. At this value, the reweighting procedure yielded a substantial reduction in *χ*^2^ while maintaining a high effective sample size (*N*_*eff*_ = 0.72), indicating that 72% of the original MD frames contribute significantly to the posterior ensemble. This high *N*_*eff*_ suggests that the initial force field sampled the relevant conformational space reasonably well, requiring only a small rebalancing of state populations rather than a selection of rare, high-energy outliers to satisfy the experimental constraints. Similar results for *θ* are seen for APO and IvA states as well (see Figures S10 and S11).

**Figure 4:**
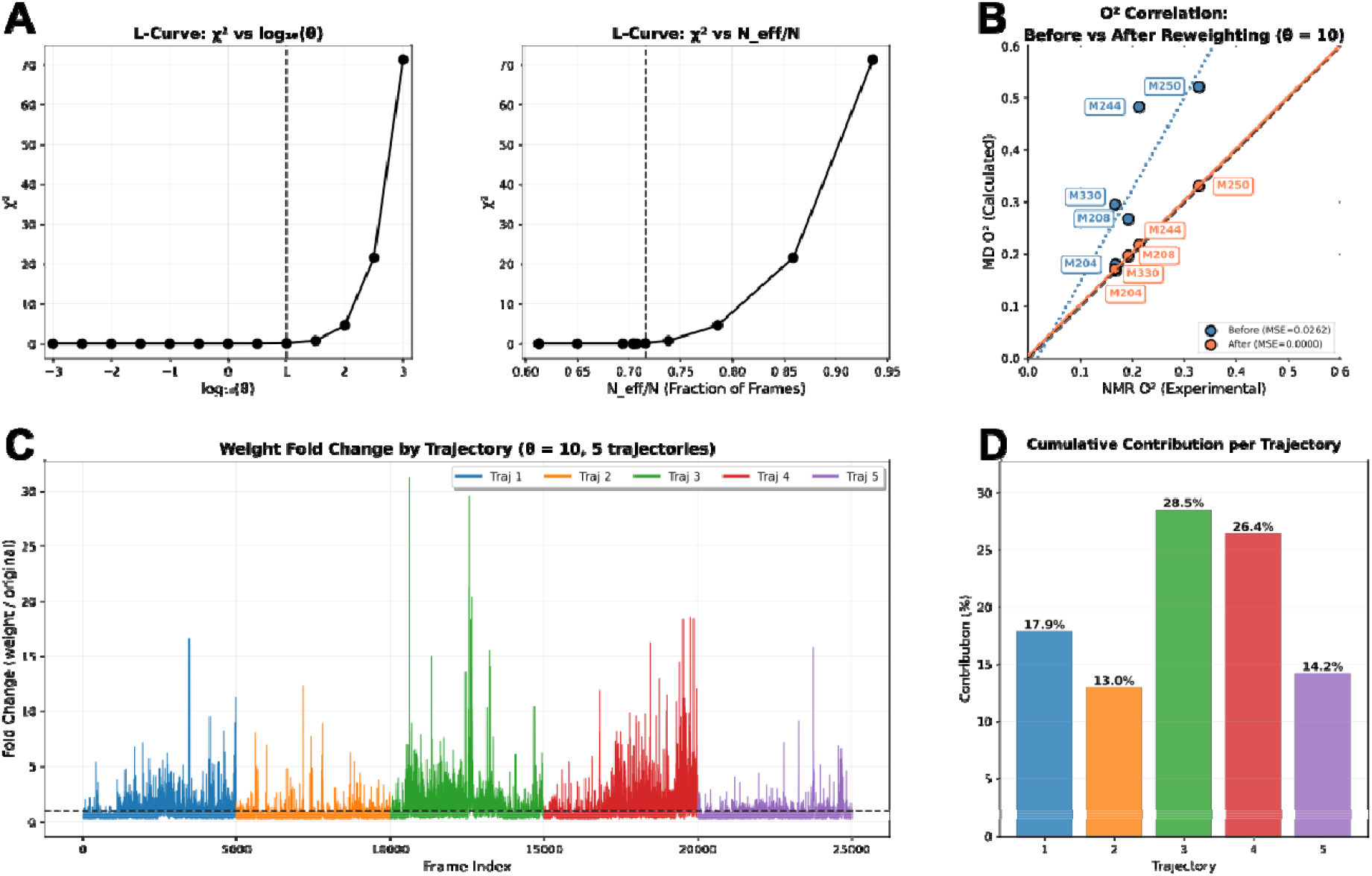
Reweighting of MD frames to match experimental NMR data. A) Two L-curve plots illustrating the relationship between the chi-squared (χ^2^) values and parameters related to model fitting. The first plot shows χ^2^ versus log_10_(θ), indicating a maximum θ value where χ^2^ remains low, suggesting an optimal range for θ. The second plot depicts χ^2^ versus effective sample size (N_eff), demonstrating that at θ = 10 (marked by a dashed vertical line on both plots), N_eff reaches a high value of 0.72, signifying a robust model fit while maintaining a high entropy of the weights. The dashed vertical lines highlight the position of θ = 10 in each plot for reference. B) Correlation plot comparing molecular dynamics (MD)-derived O^2^_axis_ values with nuclear magnetic resonance (NMR)-derived O^2^_axis_ values. The blue dots and blue dashed trend line represent the data before reweighting, whereas the orange dots and solid orange trend line correspond to the data after reweighting with θ = 10. A perfect correlation is indicated by a black-dashed line. The mean squared error (MSE) from the trend line is provided in the legend, demonstrating improved agreement and linearity after reweighting. C) Plot of fold change versus frame index across all five MD trajectories, color-coded as indicated in the legend. The dashed horizontal line at fold change = 1.0 represents no change, serving as a reference for evaluating deviations in each trajectory over time. D) Cumulative contribution of each trajectory expressed as a percentage, with all five trajectories normalized to sum to 100%. This plot quantifies the relative influence of each trajectory on the overall ensemble, highlighting its proportional contribution to the final model.

The efficacy of the reweighting protocol was evaluated by comparing the back-calculated methyl O^2^_axis_ against the experimental values derived from NMR relaxation measurements (Figure 4B, Figures S10 and S11 for APO and IvA). Prior to reweighting (blue), the correlation between simulated and experimental values exhibited significant deviations, a phenomenon often attributed to inaccuracies in methyl rotation barriers within standard force fields. Following the application of the optimized weights (orange), the ensemble showed a marked improvement in agreement with the experimental data, as evidenced by the alignment of the data points along the diagonal and the reduction in the Mean Squared Error (MSE) (Figure 4B). This refinement confirms that the reweighted ensemble provides a more accurate representation of the fast, ps-ns timescale dynamics inherent to the NTS1 receptor in this ligand-bound state. Figure 4C displays the fold change of the weights relative to the uniform prior (*w*_*j*_/*w*^0^) across the simulation frames. While spikes in weight are visible across the 5 trajectories, a histogram of fold changes in weights shows a smooth distribution, suggesting that the reweighting does not rely on a small fraction of frames and instead smoothly redistributes weight importance (Figure S12) in line with a high *N*_*eff*_. Instead, a handful of strongly upweighted frames (far right in Figure S12) push the normalized distribution’s mode slightly below 1.0, since the mean of these weights must be 1.0. The weight distributions for APO (Figure S13) and IvA (Figure S14) also show this effect, with modes below 1.0 due to the need to compensate for the far-right tails in the weight distributions. Interestingly, for APO and IvA, there is a flat region in the distributions around 1.0, suggesting that many frames aggregate into populations consistent with the NMR data.

In addition, the cumulative contribution of the five independent MD trajectories to the final reweighted ensemble (Figure 4D) demonstrates that the posterior distribution is not dominated by a single simulation replica. Instead, the final model integrates contributions from all trajectories, ensuring that the refined ensemble retains the conformational diversity needed to capture the receptor’s entropic character, although clearly trajectories 3 and 4 contributed more to matching MD to NMR. The replicate trajectories for APO and IvA are more evenly distributed (Figures S10 and Figures S11).

The integration of experimental NMR relaxation data with molecular dynamics simulations via Maximum Entropy reweighting allows for the construction of a conformational ensemble that satisfies both physical laws (via the force field) and experimental observations. The need to reweight to match the experimental O^2^_axis_ values highlights the sensitivity of methyl dynamics to subtle features of the energy landscape, such as rotameric well depths and barriers, which are often imperfectly parameterized in empirical force fields ^38^. However, the fact that a robust fit (*N*_*eff*_ =0.72) could be achieved without collapsing the ensemble suggests that the discrepancies between the raw MD and NMR data were primarily due to inaccuracies in the relative substate populations, rather than a failure to sample the relevant conformational space. By successfully refining the ensemble to match the parameters, which serve as a dynamical proxy for conformational entropy, we hope to establish a quantitative link between the microscopic fluctuations of the receptor side-chains and the global thermodynamic state. The reweighted ensemble reveals that ligand binding does not simply “lock” the receptor into a rigid state or release it into a flexible one. Rather, it redistributes the probabilities of pre-existing microstates.

### Ligand-Dependent Remodeling of the Entropic Landscape

To translate the microscopic reorientation of side-chains into thermodynamic quantities, we calculated the conformational entropy (*S*_*SC*_) for all residues possessing three or fewer *χ* angles using the reweighted probability distributions^44^. Note that calculations of S_SC_ proved intractable for residues with *χ* angles greater than 3 (Lysine and Arginine). Also, Glycine was excluded (no *χ* angle) and so was Alanine since its single *χ* angle is completely symmetric and doesn’t contribute to entropy. The total of the S_SC_ (dimensionless) values over all amino acids summed to Apo: 965.95, NT8-13: 941.03, and IvA: 966.30. The interpretation of these magnitudes suggests Apo and IvA have similar and higher global entropy than NT8-13. These results differ in interpretation from those reported by Bumbak et al., using ^13^C*ε*-methionine chemical shift-based global order parameter in which Apo is higher in dynamics than NT8-13, which in turn is higher then IvA. Entropy buildup curves for M244, M250, and M330 for Apo, NT8-13, and IvA are shown in Figure S15. Of note is how long it takes for the entropy curves of methionines to reach equilibrium. Because buildup did not typically reach equilibrium in an integer fraction of the entire MD frames (25000), only a single buildup was used per amino acid.

Figure 5 illustrates the entropic difference between the ligand-bound states and the apo receptor, providing a residue-resolved thermodynamic map of activation and inactivation.

**Figure 5:**
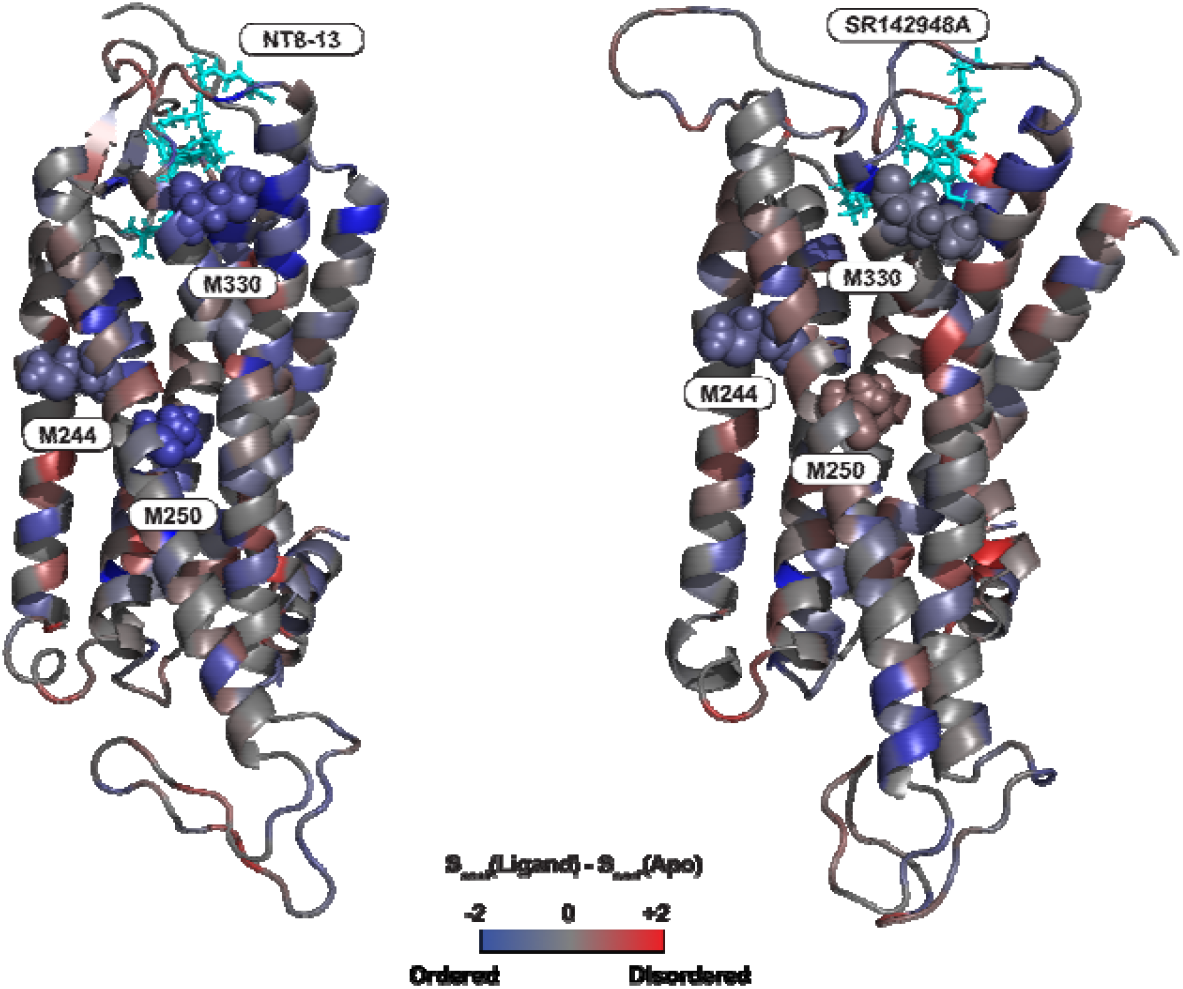
Entropic differences between APO and Ligand Bound States: Panel A shows the difference in *S*_*conf*_ calculated for every amino acid side chain with three or fewer chi angles, comparing the Apo state to the NT8-13 state. Panel B presents the same comparison but between the Apo state and IvA. Methionine side chains M244, M250, and M330 are specifically indicated. The bar gradient represents the magnitude of the difference, ranging from red (+2) through grey (0) to blue (−2).

Comparison of the agonist-bound (NT8-13) and Apo ensembles reveals a dominant trend of entropic reduction (Figure 5A, blue), consistent with the freezing out of conformational degrees of freedom upon ligand binding for M244, M250 and M330. The increase in rigidity seen for M330, in particular, is in line with our NMR result (Figure 2G). Furthermore, the orthosteric binding pocket in general undergoes significant ordering due to direct ligand contacts. Upon NT8-13 binding, the extracellular portion of TM5 is pulled inward to accommodate the ligand. This movement forces the “connector” residues into a more defined packing arrangement, thereby displacing and facilitating the outward rotation of TM6. The reduction in entropy for M244 and M250 may reflect steric constraints imposed by this tighter packing; the side chains are no longer free rotors but are “clamped” into specific orientations necessary to transmit the signal from the orthosteric pocket to transducers. In contrast, the inverse agonist (IvA) SR142948A induces less rigidity compared to the NT8-13 state (Figure 5B). Notably, M244 and M330 show minor decreases in entropy in the IvA-bound state compared to the APO state and are similar to the NT8-13 state. There is generally an increase in entropy across the region around M250.

## DISCUSSION

This study integrates advanced NMR relaxation experiments with molecular dynamics (MD) simulations, refined through Bayesian Maximum Entropy (BME) reweighting, to elucidate the fast timescale (ps–ns) side-chain dynamics and conformational entropy changes in the neurotensin receptor 1 (NTS1) upon ligand binding. The use of a minimal methionine variant of NTS1 (*enNTS*1Δ*M*4) allowed site-specific probing of local dynamics across critical receptor domains, providing a detailed thermodynamic and dynamic landscape of ligand-induced modulation.

Our NMR measurements of methyl side-chain order parameters (O^2^_axis_) revealed distinct ligand-dependent dynamic signatures. The agonist NT8-13 induces a reduction in conformational flexibility at M250^5.51^ and M330^6.57^ (Figure 2G), residues proximal to the orthosteric binding pocket and the PIF motif, consistent with a “freezing out” of conformational degrees of freedom necessary for receptor activation. This observation aligns with the structural contraction of the orthosteric pocket and stabilization of extracellular loops reported in previous crystallographic studies. Conversely, the inverse agonist SR142948A appears to increase local dynamics at M250^5.51^ and M330^6.57^ (Figure 2G), suggesting that inverse agonism involves not simply rigidification but a redistribution of dynamic states that may facilitate an expanded binding pocket conformation. These findings underscore the nuanced role of fast side-chain motions in modulating receptor function beyond static structural snapshots.

The integration of experimental data with MD simulations via BME reweighting was essential to reconcile discrepancies between raw force field predictions and NMR observables. The reweighted ensembles demonstrated improved agreement with experimental O^2^_axis_ values (Figure 4, B) while maintaining high effective sample sizes (Figure 4A), indicating that the initial force fields sampled relevant conformational space but required adjustment in their state populations to reflect experimental data. This approach highlights the sensitivity of methyl dynamics to subtle energy landscape features, such as rotameric barriers, and validates the utility of combined experimental-computational frameworks for capturing protein dynamics with atomic resolution.

Importantly, the calculated conformational entropies derived from reweighted MD ensembles provide a quantitative thermodynamic interpretation of ligand effects on NTS1. The agonist-bound state is characterized by significant entropic cost in key residues, reflecting conformational constraints necessary for signaling competence (Figure 5, left). In contrast, the inverse agonist-bound state exhibits a more entropically permissive landscape, consistent with stabilization of inactive conformations and increased side-chain flexibility (Figure 5, right). These results support a model of GPCR activation in which ligands modulate the equilibrium distribution of pre-existing microstates rather than inducing discrete conformations, emphasizing the role of dynamic allostery and conformational entropy in functional selectivity.

The findings presented here contribute to a growing body of evidence that receptor function cannot be fully understood through static structures alone. The dynamic ensemble perspective, incorporating fast side-chain motions and their thermodynamic consequences, offers a more comprehensive framework for drug discovery targeting GPCRs. By elucidating how different ligands sculpt the entropic and dynamic landscape of NTS1, this work provides mechanistic insights into biased agonism and inverse agonism, potentially guiding the design of modulators with improved efficacy and selectivity.

## ACKNOWLEDGEMENTS

The project was funded by Indiana Precision Health Initiative (J.J.Z.) and NIH grants R00GM115814 (J.J.Z.) and R35GM143054 (J.J.Z.).

## REFERENCES

1. Hauser, A.S., Attwood, M.M., Rask-Andersen, M., Schioth, H.B. & Gloriam, D.E. Trends in GPCR drug discovery: new agents, targets and indications. Nat Rev Drug Discov 16, 829–842 (2017).

2. Cebi, E. et al. Cryo-electron microscopy-based drug design. Front Mol Biosci 11, 1342179 (2024).

3. Kobilka, B.K. & Deupi, X. Conformational complexity of G-protein-coupled receptors. Trends Pharmacol Sci 28, 397–406 (2007).

4. Shimada, I., Ueda, T., Kofuku, Y., Eddy, M.T. & Wuthrich, K. GPCR drug discovery: integrating solution NMR data with crystal and cryo-EM structures. Nat Rev Drug Discov 18, 59–82 (2019).

5. Zhang, M. et al. Cryo-EM structure of an activated GPCR-G protein complex in lipid nanodiscs. Nat Struct Mol Biol 28, 258–267 (2021).

6. Krumm, B.E. et al. Neurotensin Receptor Allosterism Revealed in Complex with a Biased Allosteric Modulator. Biochemistry 62, 1233–1248 (2023).

7. Cherezov, V. et al. High-resolution crystal structure of an engineered human beta2-adrenergic G protein-coupled receptor. Science 318, 1258–65 (2007).

8. Deluigi, M. et al. Complexes of the neurotensin receptor 1 with small-molecule ligands reveal structural determinants of full, partial, and inverse agonism. Sci Adv 7 (2021).

9. Deluigi, M. et al. Crystal structure of the alpha(1B)-adrenergic receptor reveals molecular determinants of selective ligand recognition. Nat Commun 13, 382 (2022).

10. Motlagh, H.N., Wrabl, J.O., Li, J. & Hilser, V.J. The ensemble nature of allostery. Nature 508, 331–9 (2014).

11. Ziarek, J.J., Baptista, D. & Wagner, G. Recent developments in solution nuclear magnetic resonance (NMR)-based molecular biology. J Mol Med (Berl) 96, 1–8 (2018).

12. Cooper, A. & Dryden, D.T. Allostery without conformational change. A plausible model. Eur Biophys J 11, 103–9 (1984).

13. Wand, A.J. & Sharp, K.A. Measuring Entropy in Molecular Recognition by Proteins. Annu Rev Biophys 47, 41–61 (2018).

14. Lee, A.L. & Wand, A.J. Microscopic origins of entropy, heat capacity and the glass transition in proteins. Nature 411, 501–4 (2001).

15. Kasinath, V., Sharp, K.A. & Wand, A.J. Microscopic insights into the NMR relaxation-based protein conformational entropy meter. J Am Chem Soc 135, 15092–100 (2013).

16. Ramirez-Virella, J. & Leinninger, G.M. The Role of Central Neurotensin in Regulating Feeding and Body Weight. Endocrinology 162 (2021/05/01).

17. White, J.F. et al. Structure of the agonist-bound neurotensin receptor. Nature 490, 508–13 (2012).

18. Bumbak, F. et al. Stabilization of pre-existing neurotensin receptor conformational states by beta-arrestin-1 and the biased allosteric modulator ML314. Nat Commun 14, 3328 (2023).

19. Barak, L.S. et al. ML314: A Biased Neurotensin Receptor Ligand for Methamphetamine Abuse. ACS Chem Biol 11, 1880–90 (2016).

20. Clark, L.D. et al. Ligand modulation of sidechain dynamics in a wild-type human GPCR. Elife 6(2017).

21. Baumann, C. et al. Side-chain dynamics of the alpha(1B)-adrenergic receptor determined by NMR via methyl relaxation. Protein Sci 32, e4801 (2023).

22. Kofuku, Y. et al. Functional dynamics of deuterated beta2-adrenergic receptor in lipid bilayers revealed by NMR spectroscopy. Angew Chem Int Ed Engl 53, 13376–9 (2014).

23. Xu, J. et al. Conformational Complexity and Dynamics in a Muscarinic Receptor Revealed by NMR Spectroscopy. Mol Cell 75, 53–65 e7 (2019).

24. Bumbak, F. et al. Optimization and (13)CH(3) methionine labeling of a signaling competent neurotensin receptor 1 variant for NMR studies. Biochim Biophys Acta Biomembr 1860, 1372–1383 (2018).

25. Lipari, G. & Szabo, A. Model-free approach to the interpretation of nuclear magnetic resonance relaxation in macromolecules. 1. Theory and range of validity. Journal of the American Chemical Society 104, 4546–4559 (1982).

26. Lipari, G. & Szabo, A. Model-free approach to the interpretation of nuclear magnetic resonance relaxation in macromolecules. 2. Analysis of experimental results. Journal of the American Chemical Society 104, 4559–4570 (1982).

27. Krishnan, M. & Smith, J.C. Reconstruction of protein side-chain conformational free energy surfaces from NMR-derived methyl axis order parameters. J Phys Chem B 116, 4124–33 (2012).

28. O’Brien, E.S. et al. Membrane Proteins Have Distinct Fast Internal Motion and Residual Conformational Entropy. Angew Chem Int Ed Engl 59, 11108–11114 (2020).

29. Sun, H., Kay, L.E. & Tugarinov, V. An optimized relaxation-based coherence transfer NMR experiment for the measurement of side-chain order in methyl-protonated, highly deuterated proteins. J Phys Chem B 115, 14878–84 (2011).

30. Bottaro, S., Bengtsen, T. & Lindorff-Larsen, K. Integrating Molecular Simulation and Experimental Data: A Bayesian/Maximum Entropy Reweighting Approach. Methods Mol Biol 2112, 219–240 (2020).

31. Bumbak, F. et al. Ligands selectively tune the local and global motions of neurotensin receptor 1 (NTS(1)). Cell Rep 42, 112015 (2023).

32. Hyberts, S.G., Milbradt, A.G., Wagner, A.B., Arthanari, H. & Wagner, G. Application of iterative soft thresholding for fast reconstruction of NMR data non-uniformly sampled with multidimensional Poisson Gap scheduling. J Biomol NMR 52, 315–27 (2012).

33. Delaglio, F. et al. NMRPipe: a multidimensional spectral processing system based on UNIX pipes. J Biomol NMR 6, 277–93 (1995).

34. Hardy, R. & Cottington, R.L. Viscosity of deuterium oxide and water in the range 5 to 125 C. Journal of research of the National Bureau of Standards 42(1949).

35. Jo, S., Kim, T., Iyer, V.G. & Im, W. CHARMM-GUI: a web-based graphical user interface for CHARMM. J Comput Chem 29, 1859–65 (2008).

36. Cheng, X., Jo, S., Lee, H.S., Klauda, J.B. & Im, W. CHARMM-GUI micelle builder for pure/mixed micelle and protein/micelle complex systems. J Chem Inf Model 53, 2171–80 (2013).

37. Best, R.B. et al. Optimization of the additive CHARMM all-atom protein force field targeting improved sampling of the backbone phi, psi and side-chain chi(1) and chi(2) dihedral angles. J Chem Theory Comput 8, 3257–3273 (2012).

38. Hoffmann, F., Mulder, F.A.A. & Schafer, L.V. Predicting NMR relaxation of proteins from molecular dynamics simulations with accurate methyl rotation barriers. J Chem Phys 152, 084102 (2020).

39. Bower, J.B. et al. Stabilization versus flexibility: detergent-dependent trade-offs in neurotensin receptor 1 GPCR ensembles. bioRxiv (2025).

40. Cesari, A. et al. Using the Maximum Entropy Principle to Combine Simulations and Solution Experiments. Computation 2018, Vol. 6, 6(2018-02-06).

41. Boomsma, W., Ferkinghoff-Borg, J. & Lindorff-Larsen, K. Combining experiments and simulations using the maximum entropy principle. PLoS Comput Biol 10, e1003406 (2014).

42. Virtanen, P. et al. SciPy 1.0: fundamental algorithms for scientific computing in Python. Nat Methods 17, 261–272 (2020).

43. Bhattacharya, S. et al. Conformational dynamics and multimodal interaction of Paxillin with the focal adhesion targeting domain. Sci Adv 11, eadt9936 (2025).

44. Hoffmann, F., Mulder, F.A.A. & Schafer, L.V. How Much Entropy Is Contained in NMR Relaxation Parameters? J Phys Chem B 126, 54–68 (2022).

45. Tugarinov, V., Sprangers, R. & Kay, L.E. Probing side-chain dynamics in the proteasome by relaxation violated coherence transfer NMR spectroscopy. J Am Chem Soc 129, 1743–50 (2007).

46. Abril-Pla, O. et al. PyMC: a modern, and comprehensive probabilistic programming framework in Python. PeerJ Comput Sci 9, e1516 (2023).

