## Supplementary Figures for "Mapping the Ligand-dependent Remodeling of the Conformational Entropy Landscape in Neurotensin Receptor 1 by NMR-guided Molecular Simulations"

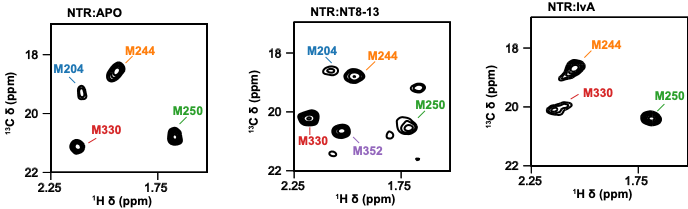

Figure S1: First SQ plane from the interleaved Triple Quantum relaxation experiment with methionine assignments indicated.

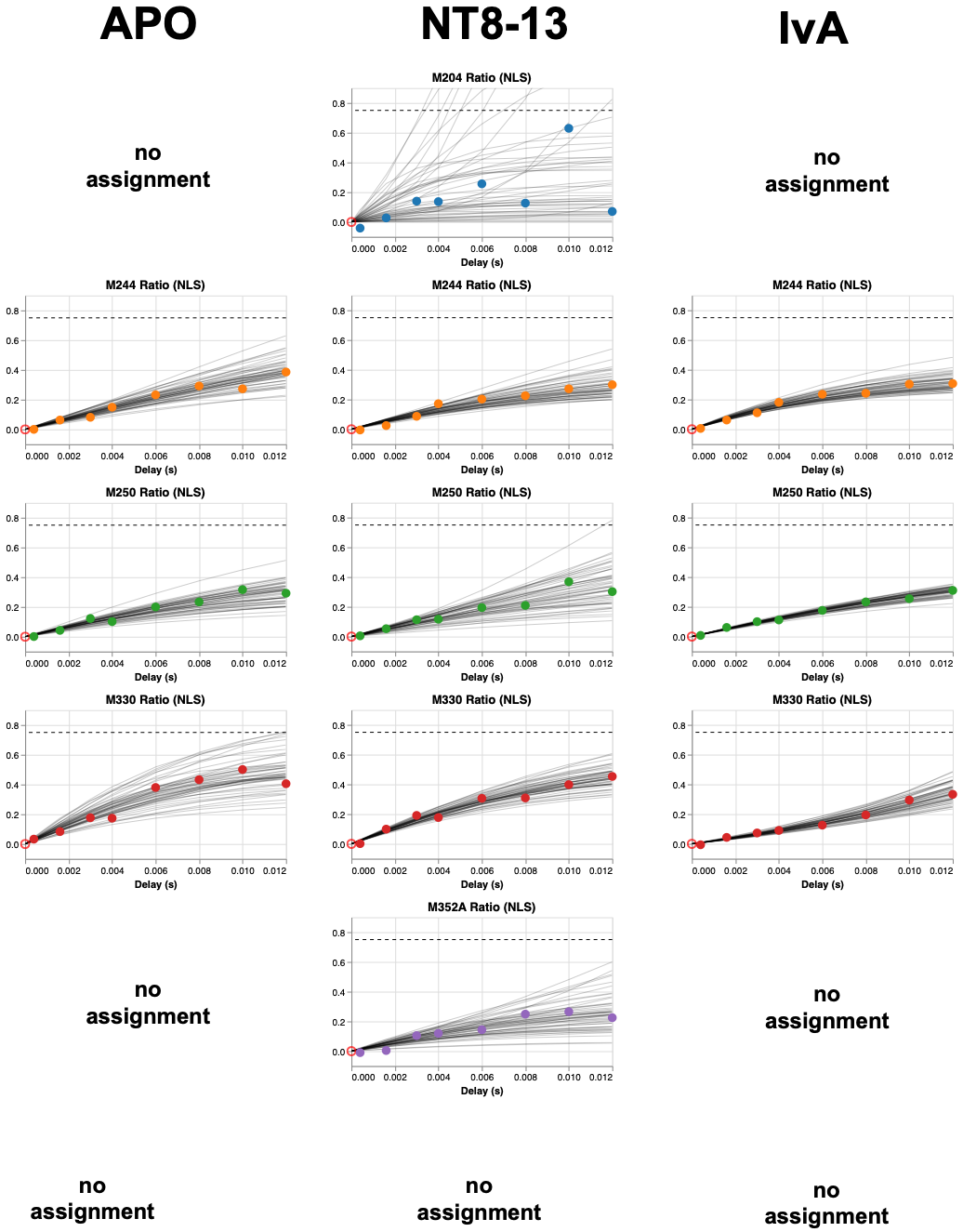

Figure S2: Model fitting of the time series ratios in the 3Q relaxation experiment using Eq. 1. High errors were seen, motivating another approach.

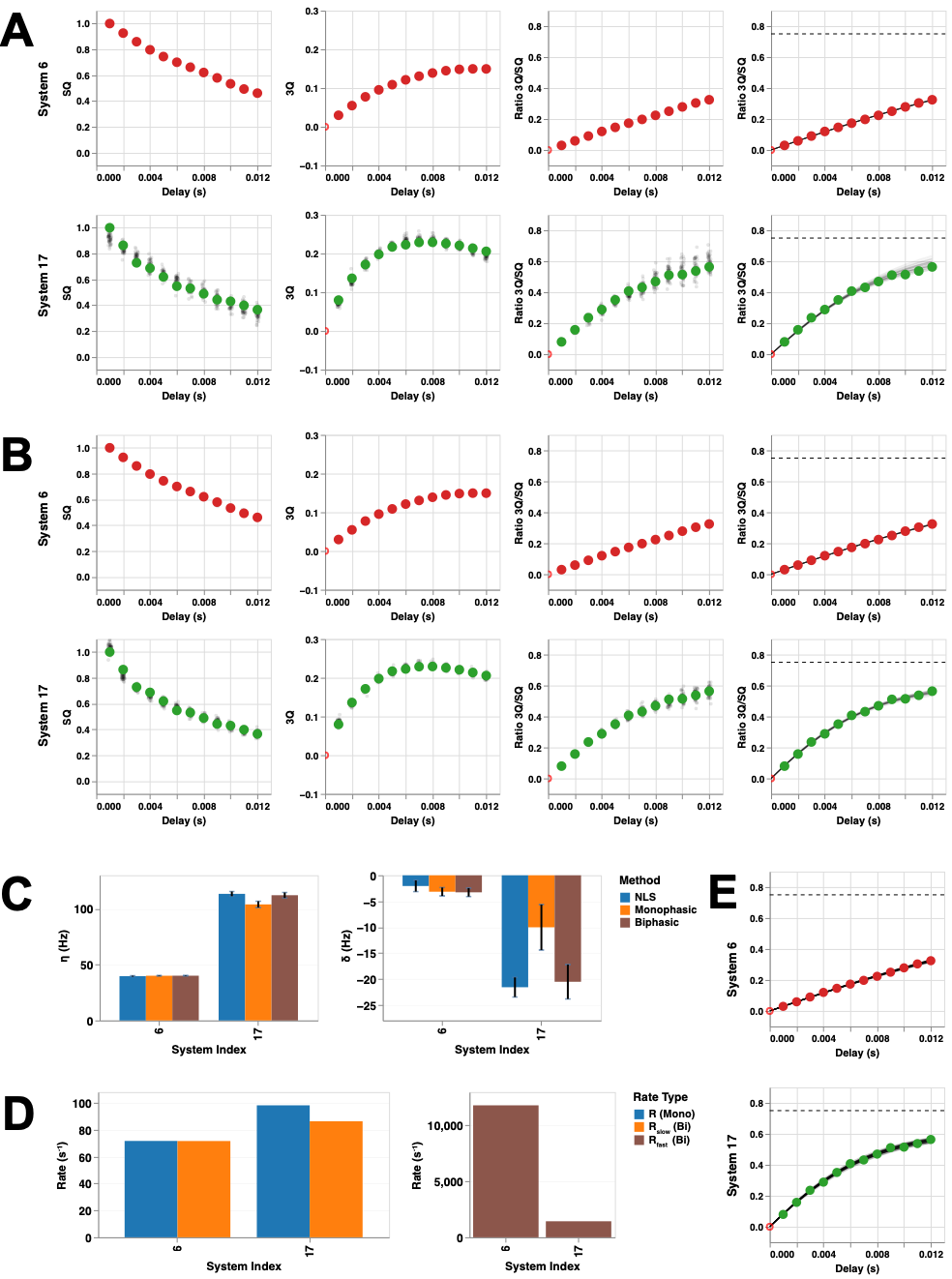

Figure S3: Fitting of two (Panel A and B) example systems from Ubiquitin α7α7 ( τ_c_ ~ 6 ns) using the approach of Eq. 2 and 3. Panel C shows the agreement between using Eq. 1 with nonlinear least squares (NLS) and our approach, which includes either monophasic or biphasic relaxation terms. In Panel D, we show that system 6 produces a very large second relaxation rate (brown), which therefore contributes little to the fit. Panel E is a fit of the data from traditional NLS.

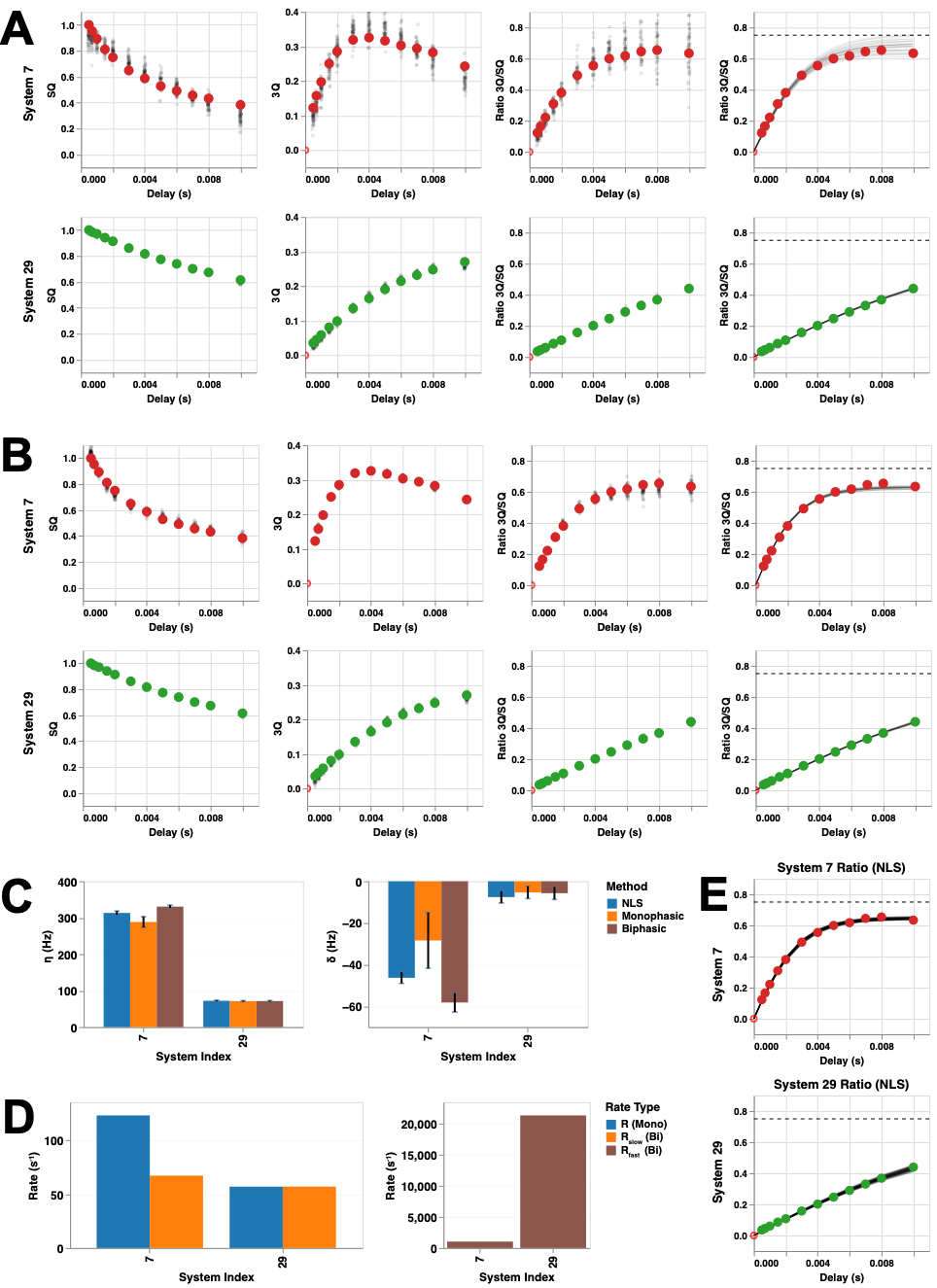

Figure S4: Fitting of two (Panel A and B) example systems from α7α7 ( τ_c_ ~ 107 ns) using the approach of Eq. 2 and 3. Panel C shows the agreement between using Eq. 1 with nonlinear least squares (NLS) and our approach, which includes either Monophasic or Biphasic relaxation terms. In Panel D, we show that system 29 produces a very large second relaxation rate (brown), which doesn’t contribute to the fit. System 7, however, has a significant second rate (brown) compared to the first (orange). The effect of this can be seen by comparing Panel A and B, with B (two relaxation terms) fitting the data much better.

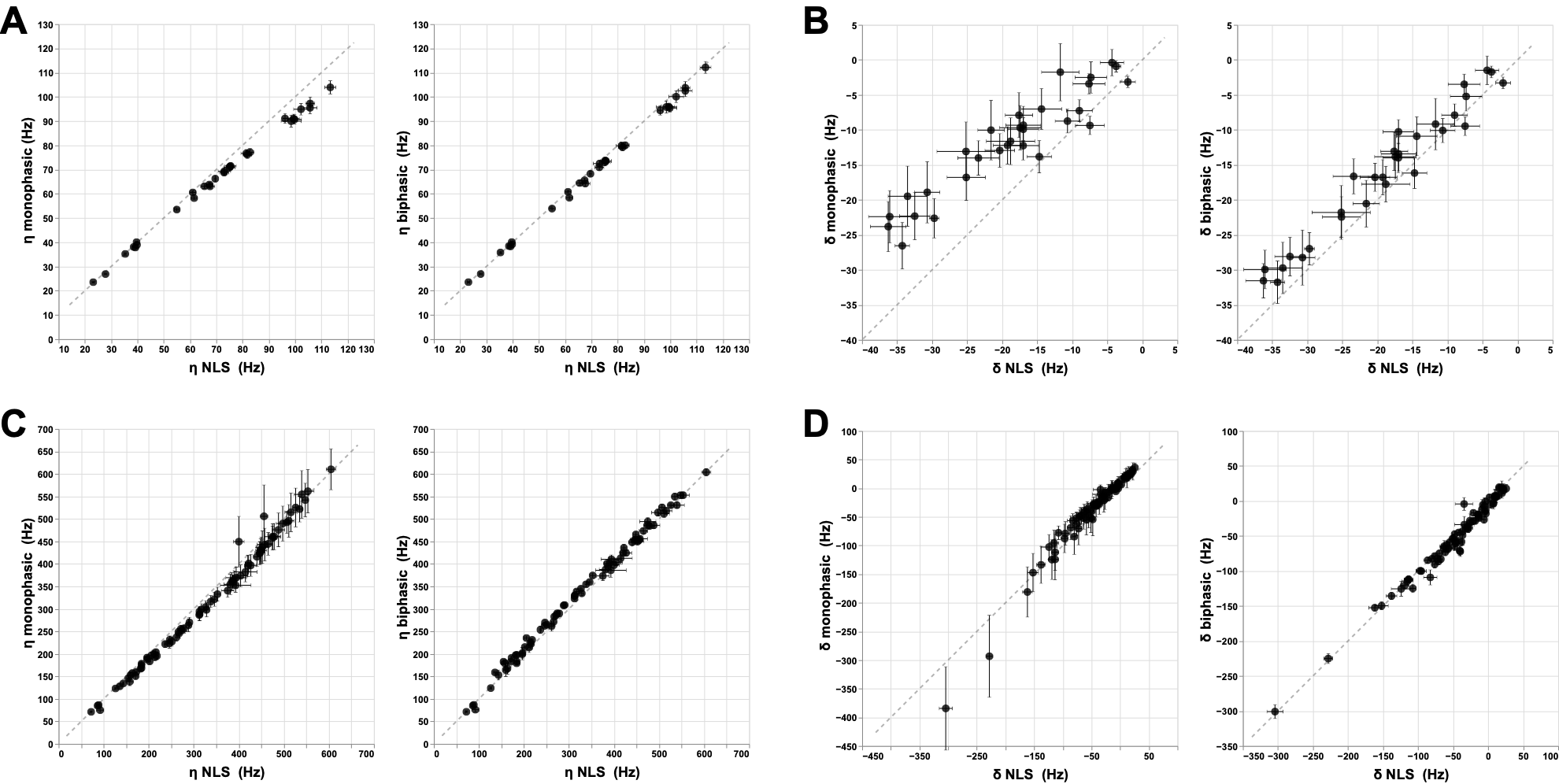

Figure S5: Correlation plots of η and δ values extract by the monophasic or biphasic method compared to NLS for all assigned systems in Ubiquitin and α7α7. Panel A (left) shows the correlation for Ubiquitin η using a monophasic model compared to the biphasic model (right). Panel B is the same but for δ values. Panel C (left) ) shows the correlation for α7α7 η using a monophasic model compared to the biphasic model (right). Panel D is the same but for δ values. In each case, using the biphasic model improves the correlation with example (high quality) data from Vitali Tugarinov and the correlation is high.

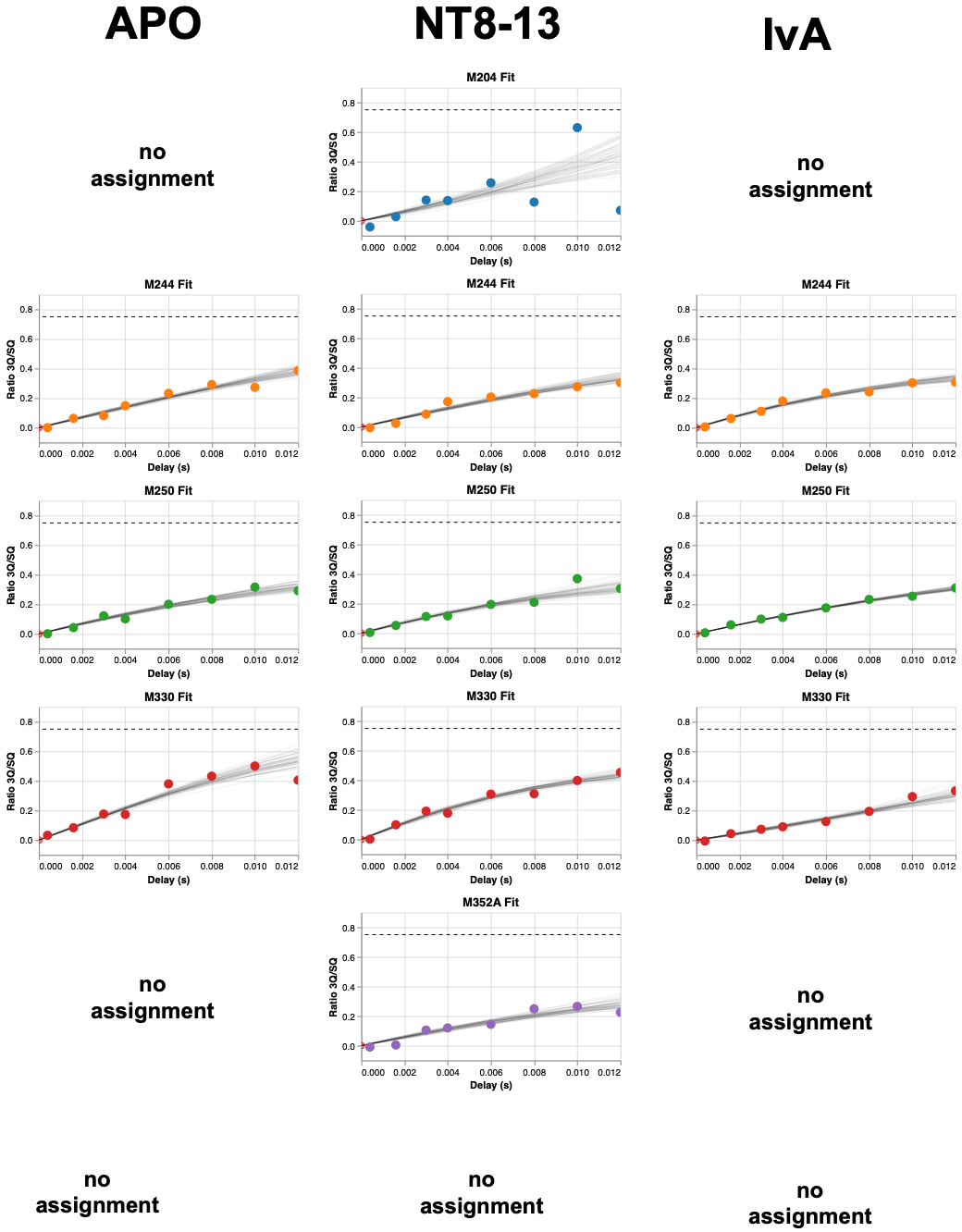

Figure S6: Model fitting of the time series ratios in the 3Q relaxation experiment using the biphasic approach of Eq. 2 and 3. There is a significant improvement in model fit error compared to Figure S2.

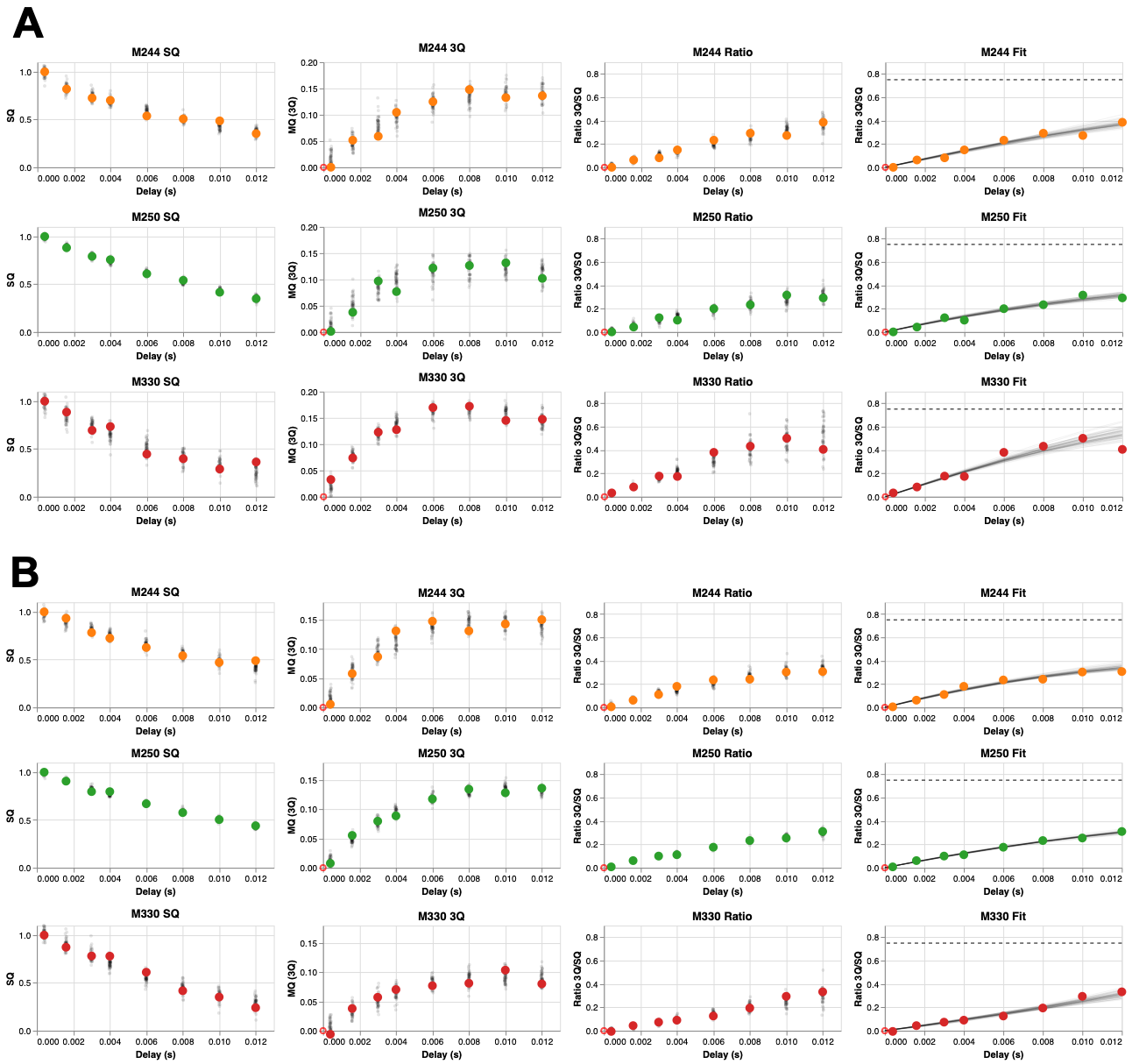

Figure S7: Panel A: Complete system fits for the APO state for Eq. 2 and 3 and their ratio, along with the final fitted model. Panel B: The same as Panel A but for the IvA state.

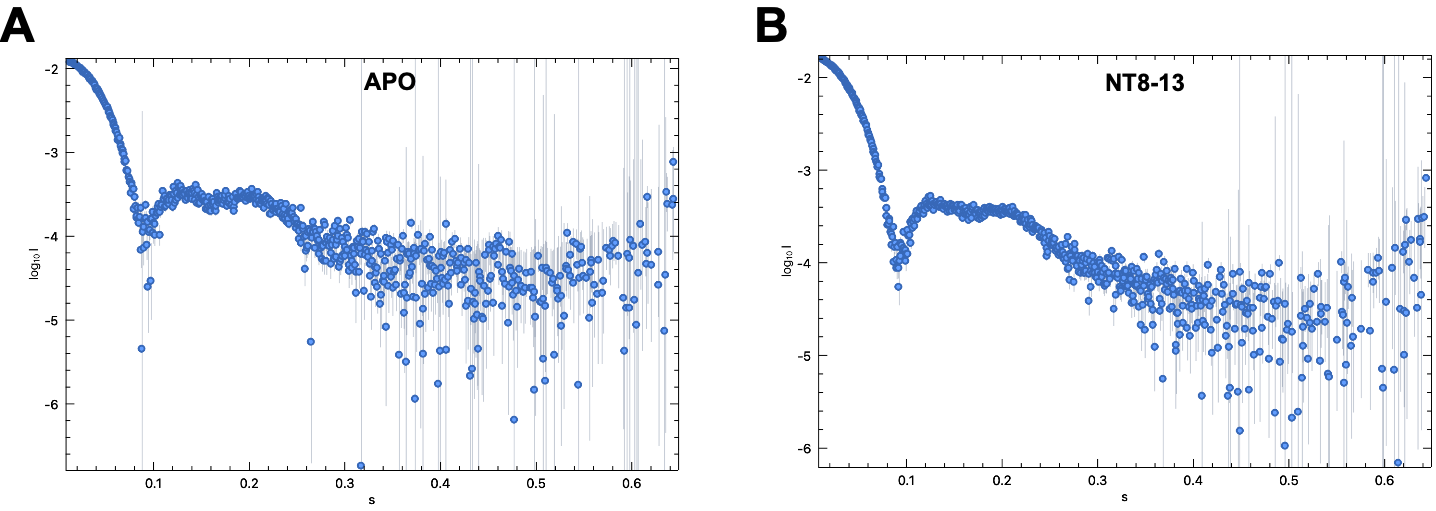

Figure S8: Raw scattering data from SEC-SAX, demonstrating a similar particle shape and size for the APO and NT8-13 state.

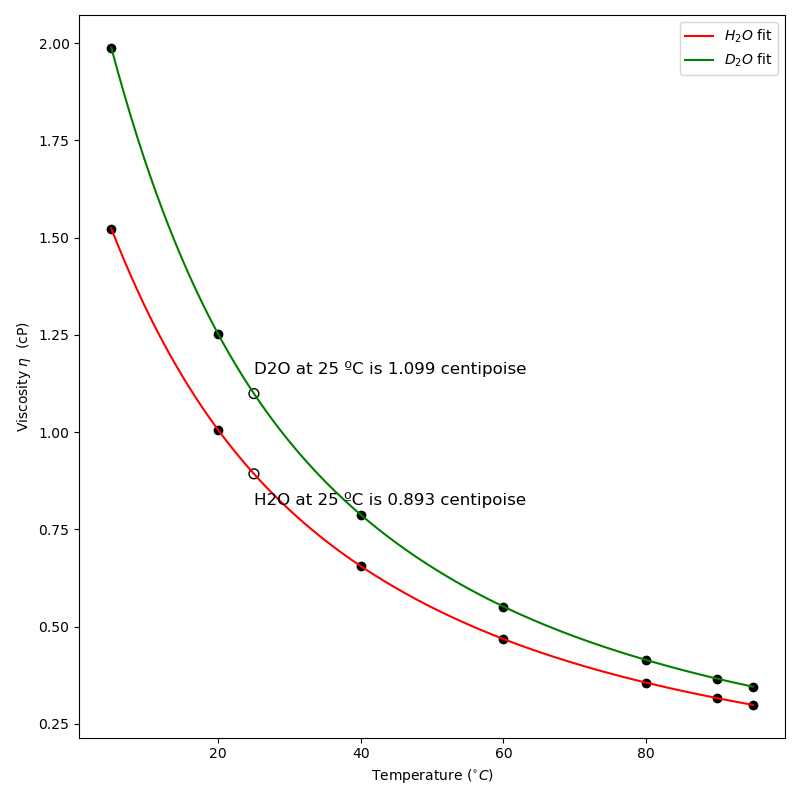

Figure S9: Calibration curve of D2O viscosity from data taken from Hardy et. al. Calibration curve was fitted to data points using a third order polynomial.

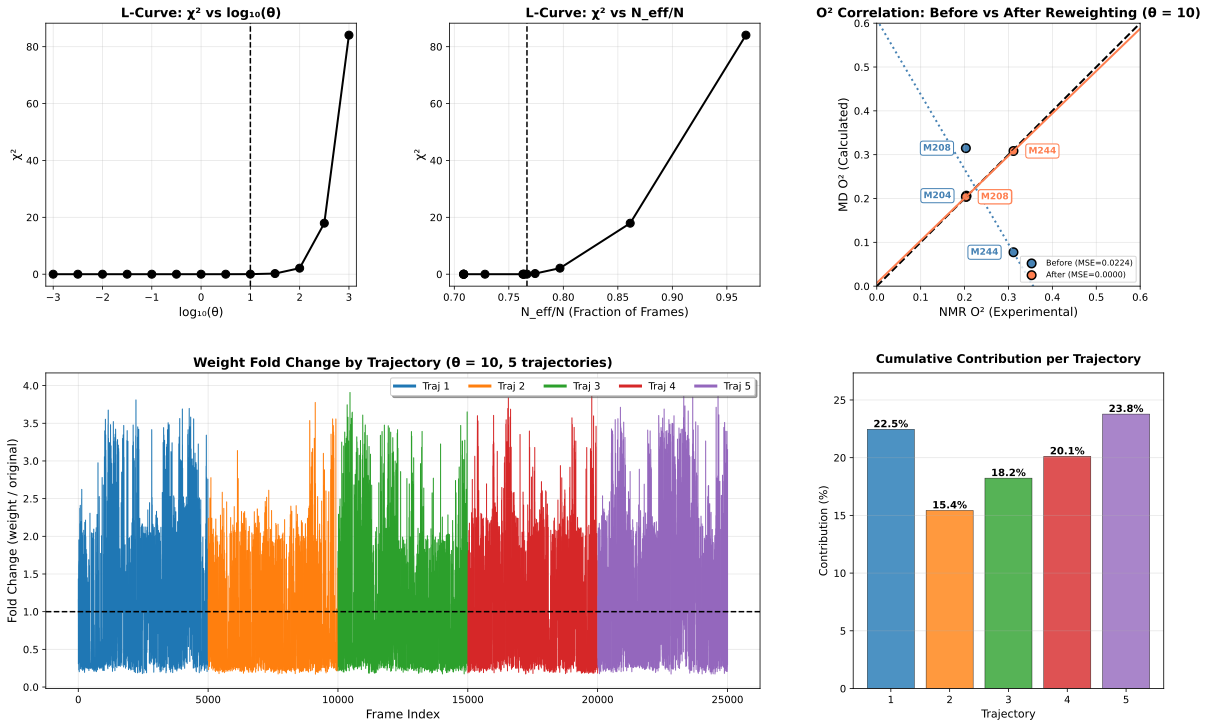

Figure S10: Same plot as Figure 4, but for the APO state.

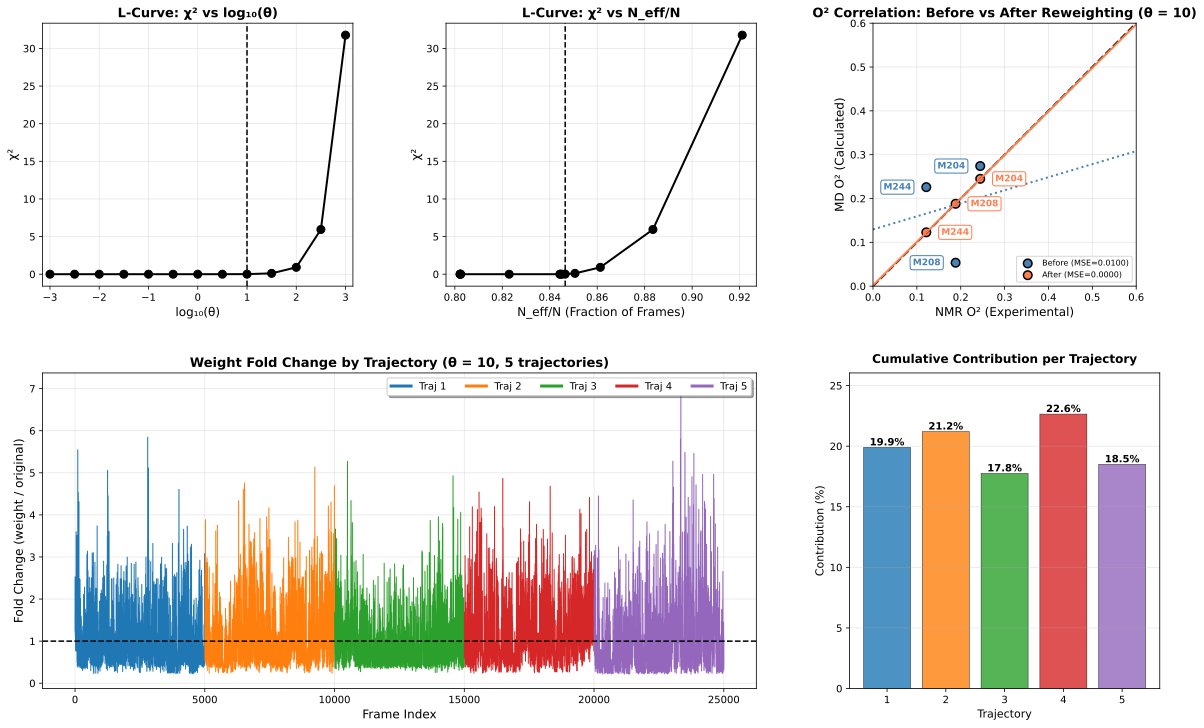

Figure S11: Same plot as Figure 4, but for the IvA state.

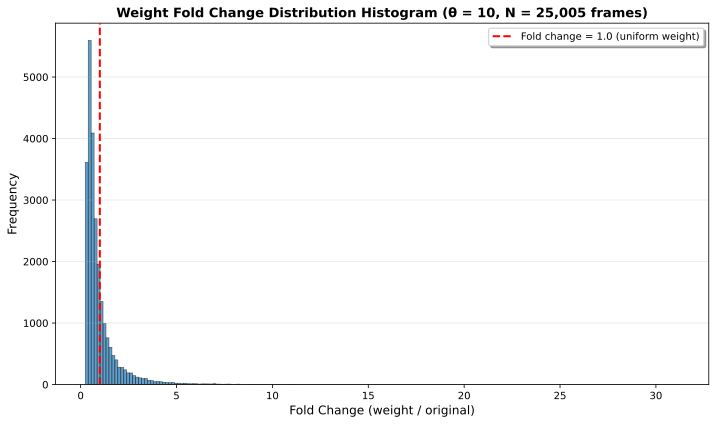

Figure S12: Fold change in weight distribution for NT8-13. The presence of a heavy tail to the right pushes the mode to under 1.0, however the distribution is smooth suggesting there is not a simple subpopulation of ‘correct’ frames.

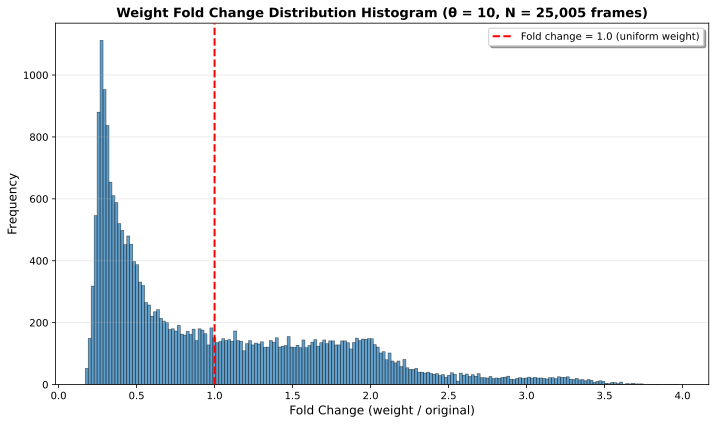

Figure S13: Fold change in weight distribution for APO. An extended population of frames hover around 1.0, but the existence of a right-side tail pushes the mode to below 1.0.

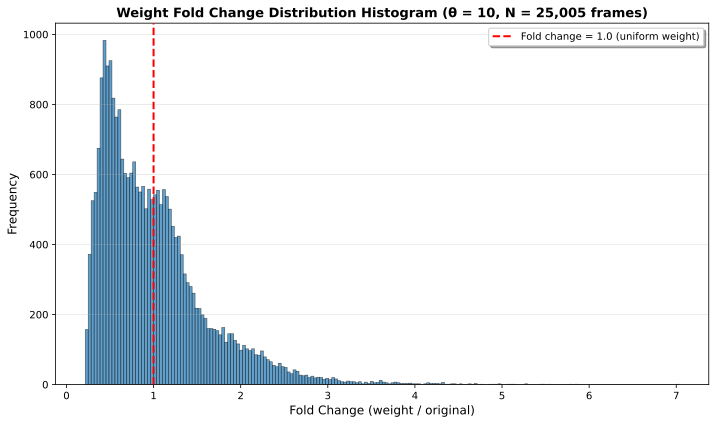

Figure S14: Fold change in weight distribution for IvA. Again, there is a population of frames hover around 1.0, but the existence of a right-side tail pushes the mode to below 1.0.

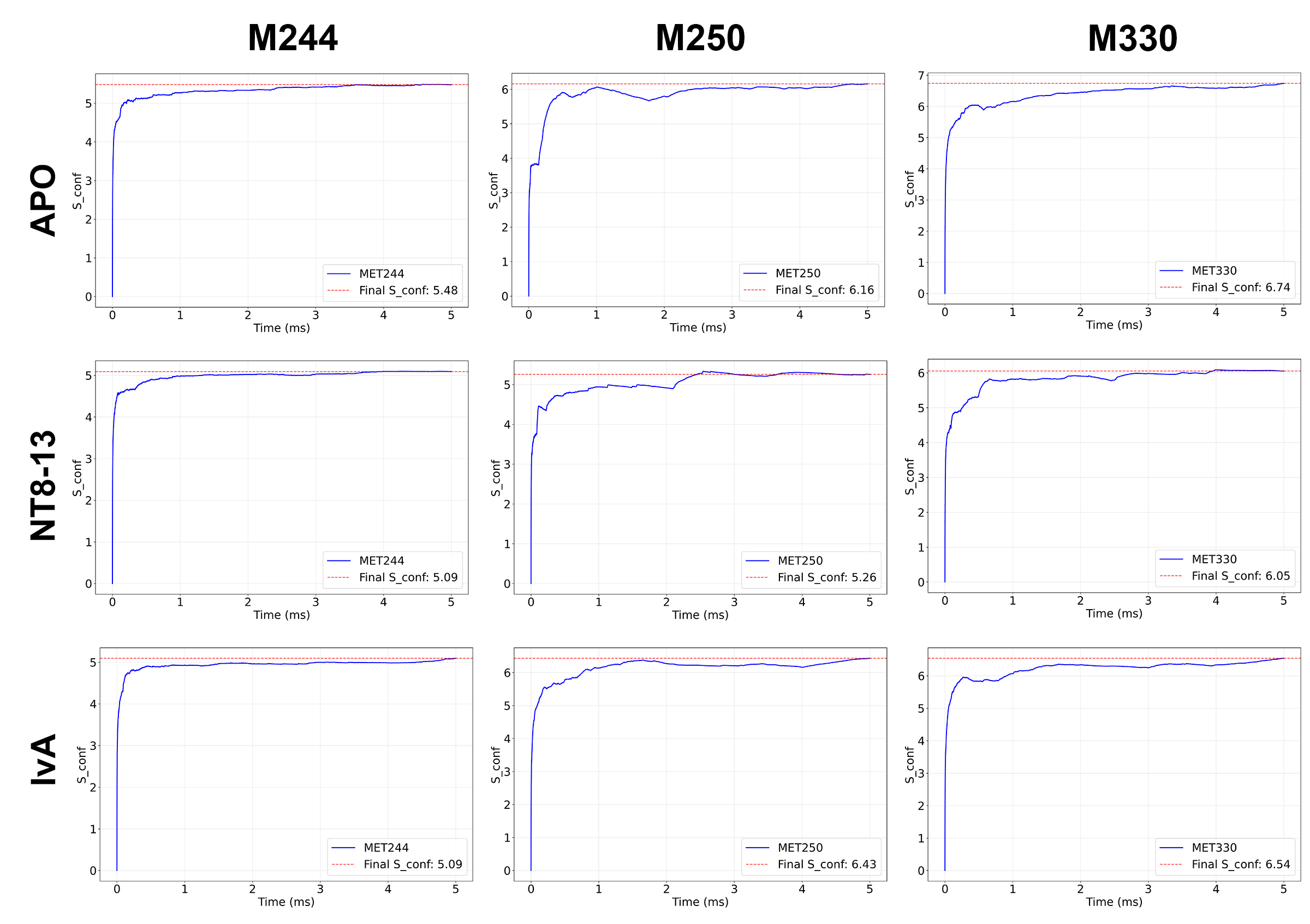

Figure S15: Entropic Buildup During MD Simulations for M244, M250 and M330 (columns) for APO, NT8-13 and IvA (rows):
